# Multiplexed live-cell imaging for drug responses in patient-derived organoid models of cancer

**DOI:** 10.1101/2023.11.15.567243

**Authors:** Kaitriana E. Colling, Emily L. Symons, Lorenzo Buroni, Hiruni K. Sumanisiri, Jessica Andrew-Udoh, Emily Witt, Haley A. Losh, Abigail M. Morrison, Kimberly K. Leslie, Christopher J. Dunnill, Johann S. De Bono, Kristina W. Thiel

## Abstract

Patient-derived organoid (PDO) models of cancer are a multifunctional research system that better recapitulates human disease as compared to cancer cell lines. PDO models can be generated by culturing patient tumor cells in extracellular basement membrane extracts (BME) and plating as three-dimensional domes. However, commercially available reagents that have been optimized for phenotypic assays in monolayer cultures often are not compatible with BME. Herein we describe a method to plate PDO models and assess drug effects using an automated live-cell imaging system. In addition, we apply fluorescent dyes that are compatible with kinetic measurements to simultaneously quantitate cell health and apoptosis. Image capture can be customized to occur at regular time intervals over several days. Users can analyze drug effects in individual Z-plane images or a Z Projection of serial images from multiple focal planes. Using masking, specific parameters of interest are calculated, such as PDO number, area, and fluorescence intensity. We provide proof-of-concept data demonstrating the effect of cytotoxic agents on cell health, apoptosis and viability. This automated kinetic imaging platform can be expanded to other phenotypic readouts to understand diverse therapeutic effects in PDO models of cancer.

**SUMMARY:** Patient-derived tumor organoids are a sophisticated model system for basic and translational research. This methods article details the use of multiplexed fluorescent live-cell imaging for simultaneous kinetic assessment of different organoid phenotypes.

## INTRODUCTION

Patient-derived tumor organoids (PDOs) are rapidly emerging as a robust model system to study cancer development and therapeutic responses. PDOs are three-dimensional (3D) cell culture systems that recapitulate the complex genomic profile and architecture of the primary tumor^1,2.^ Unlike traditional two-dimensional (2D) culture of immortalized cancer cell lines, PDOs capture and maintain intratumoral heterogeneity^3,4,^making them a valuable tool for both mechanistic and translational research. Although PDOs are becoming an increasingly popular model system, commercially available reagents and analysis methods for cellular effects that are compatible with PDO cultures are limited.

The lack of robust methods to analyze subtle changes in treatment response hinders clinical translation. The gold standard cell health reagent in 3D cultures, CellTiter-Glo 3D, utilizes ATP levels as a determinant of cell viability^5,6^. While this reagent is a useful for endpoint assays, there are several caveats, most notably the inability to use samples for other purposes after completion of the assay.

Live-cell imaging is a sophisticated form of kinetic microscopy that, when combined with fluorescent reagents, has the capacity to quantify a variety of cell health readouts within PDO models, including apoptosis^7-9^ and cytotoxicity^10^. Indeed, live-cell imaging has been integral to high throughput screening of compounds in 2D platforms^11,12.^Systems such as the Incucyte have made the technology affordable and thus accessible to research groups in a variety of settings. However, application of these systems to analyze 3D cultures is still in its infancy.

Herein we describe a method to assess drug response in PDO models of cancer using multiplexed live-cell imaging (**Figure 1**). Through analysis of Bright Field images, changes in organoid size and morphology can be kinetically monitored. Furthermore, cellular processes can be simultaneously quantified over time using fluorescent reagents, such as Annexin V Red Dye for apoptosis and Cytotox Green Dye for cytotoxicity. The methods presented are optimized for the Cytation 5 live-cell imaging system, but this protocol may be adapted across different live-cell imaging platforms.

**Figure 1.**
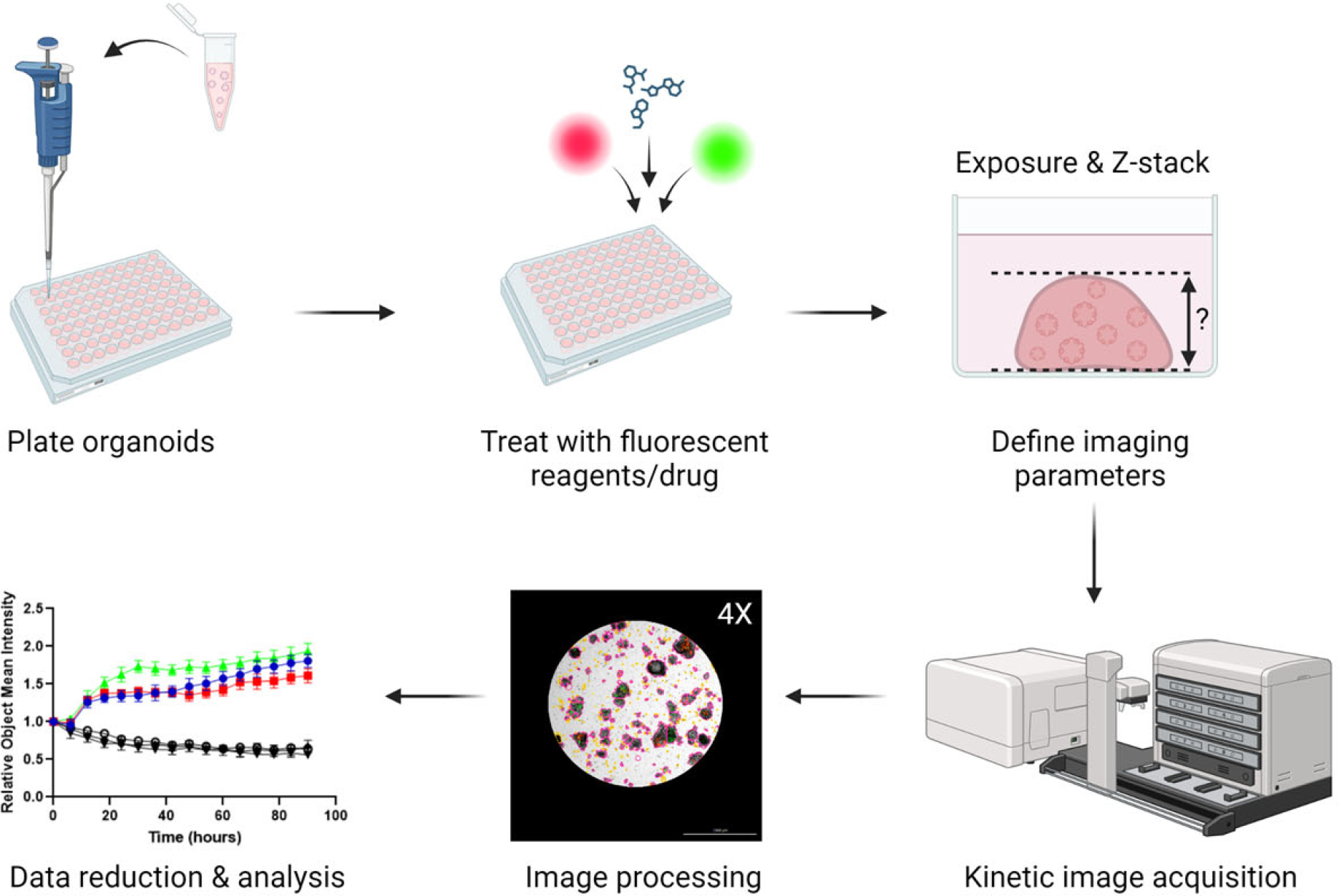
Overview of plating, imaging and analysis protocol. PDOs are plated in a 96-well plate and treated with fluorescent dyes and drugs. Imaging parameters for the experiment (e.g., Exposure, Z-stack) are created in the Gen5 software. Images are acquired by the Cytation 5 and processed in Gen5, and data are exported for further analysis.

## PROTOCOL

### Ethics Statement

Studies using human tumor specimens were reviewed and approved by the University of Iowa Institutional Review Board (IRB), protocol #201809807 and performed in accordance with the ethical standards as laid down in the 1964 Helsinki Declaration and its later amendments. Informed consent was obtained from all subjects participating in the study. Inclusion criteria include a diagnosis of cancer and availability of tumor specimen.

### Plating intact PDOs in a 96-well plate

1. Prepare reagents:
  1. Preheat 96-well plates at 37 °C overnight and thaw BME overnight at 4 °C.
  2. Prepare full Organoid Culture Media optimized for culturing the cancer type of interest. Specific culture media used for experiments shown herein are provided in **Supplemental Table S1**.
    1. *Note: Media components may need to be modified for different tumor types. For example, the Organoid Culture Media is supplemented with 100 nM estradiol for gynecologic tumors* ^13^.
    2. *Note: Prepared media is stable at 4 °C for 1 month. For long term storage, aliquot into 50 mL tubes and store at -20 °C*.
2. Prepare two separate aliquots of Organoid Culture Media at 4 °C and 37 °C.
  1. For example, if 60 wells are being plated in a 96-well plate, 6 mL of warm Organoid Culture Media and 150 µL of ice-cold Organoid Culture Media is required.
3. Prepare Organoid Wash Buffer: 1X PBS supplemented with 10 mM HEPES, 1X Glutamax, 5 mM EDTA, and 10 µM Y-27632. Store at 4 °C
4. Harvest PDOs cultured in a 24-well plate: All steps should be performed on ice or at 4 °C unless otherwise noted.
  1. Aspirate media from each well using a vacuum line system.
  2. Add 500 µL ice-cold Organoid Wash Buffer and gently pipette 2-3X using a 1000 µL pipettor.
  3. Incubate plate on ice for 10 minutes.
  4. Transfer the contents of each well to a 50 mL conical tube. To ensure that all the PDOs are in suspension, rinse each well with an additional 300 µL of Organoid Wash Buffer and transfer to the 50 mL conical tube.
  5. Centrifuge for 5 minutes at 350 x g at 4 °C.
  6. Aspirate supernatant from BME/organoid pellet using a vacuum line system, leaving ∼ 5 mL remaining in tube. Add an additional 20 mL of Organoid Wash Buffer and gently resuspend the pellet using a 10 mL serological pipette.
  7. Incubate on ice for 10 minutes.
  8. Centrifuge for 5 minutes at 350 x g at 4 °C.
  9. Aspirate the supernatant with vacuum line system, taking care not to disrupt the PDO pellet.
5. Plating PDOs in a 96-well plate: All steps should be performed on ice unless otherwise noted.
  1. Resuspend PDO pellet in appropriate amount of ice-cold Organoid Culture Media to create a PDO suspension.
    1. *Note: To calculate the amount of Organoid Culture Media, determine the number of wells to be plated in a 96-well plate, taking into consideration that PDOs are plated in a 5 µL dome in a 1:1 ratio of Organoid Culture Media and BME. For example, when plating one 96-well plate and using only the inner 60 wells, the total amount of PDO suspension needed will be 300 µL: 150 µL Organoid Culture Media and 150 µL BME*.
    2. *Note: For models that exhibit optimal growth at different percentages of BME, the ratio of BME:media may be modified in this step, though it is important to standardize the ratio across all assays for each specific model. To account for pipetting error, add 10% volume to each component*.
  2. Count the number of PDOs: Transfer 2.5 µL PDO suspension to an ice-cold 1.5 mL Microcentrifuge tube and mix with 2.5 µL BME. Transfer the 5 µL mixture onto a clean glass microscope slide. Do not coverslip the slide. The mixture will solidify into a dome. Visualize using a Bright Field microscope at 4X. Count the number of PDOs in the 5 µL mixture; the goal is to have roughly 25-50 PDOs per 5 µL dome.
    1. *Note: If the desired density is not achieved in the test mixture, adjust the final volume of the PDO suspension either by adding more Organoid Culture Media or centrifuging the PDO suspension and resuspending the PDO pellet in a lower volume of ice-cold Organoid Culture Media. Regardless of how the PDO suspension is modified in this step, the final ratio of BME:PDO suspension in Step 3 should be 1:1*.
  3. Using a 200 µL pipettor with wide bore tips, carefully mix PDO suspension with an equal amount of BME to achieve a 1:1 ratio of Organoid Culture Media to BME. Avoid bubbles, which will disrupt the integrity of the domes. Transfer contents of 15 mL conical tube to an ice-cold 1.5 mL microcentrifuge tube for easier handling in subsequent steps.
  4. Using a 20 µL pipettor, seed 5 µL domes into the center of each well of a prewarmed 96-well plate, seeding only the inner 60 wells. To ensure equal distribution of the PDOs, periodically gently pipet the contents of the 1.5 mL tube with a 200 µL pipettor with wide-bore tips.
  5. After all wells have been seeded, place the lid on the plate and gently invert. Incubate the inverted plate at 37 °C for 20 minutes in the tissue culture incubator to allow domes to solidify.
    1. *Note: Inverting the plate ensures that the BME/Organoid Culture Media dome retains the 3D structure to provide adequate room for PDO formation*.
  6. Flip the plate so that it is sitting with the lid up and incubate for an additional 5 minutes at 37 °C.

### Treatments and addition of fluorescent dyes for multiplexing

1. While BME domes are solidifying in the 96-well plates, prepare dilutions of fluorescent live-cell imaging reagents. Herein we give specific parameters for multiplexing Annexin V Red Dye and Cytotox Green Dye.
2. Fluorescent reagent preparation (Day -1): Calculate the appropriate volume of Organoid Culture Media based on the number of wells to be treated, assuming each well will be treated with 100 µL of dye-dosed media. Dilute dye in pre-warmed Organoid Culture Media to the desired concentration.
  1. *Note: The total amount of media needed will vary depending on the experiment. Add 10% to the final volume to account for pipetting error. For example, to treat the inner 60 wells of a 96-well plate, prepare 6*.*6 mL of dye-dosed media (****Table 1***).

**Table 1:**
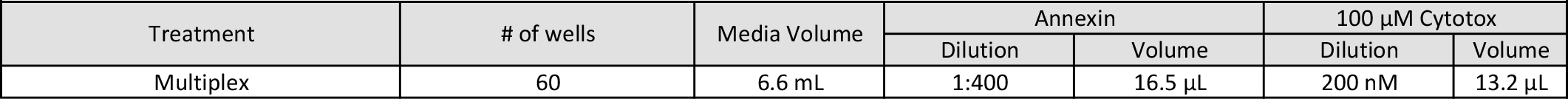
Example Multiplexing Experiment. Annexin V binds exposed phosphatidyl serine on the outer leaflet of apoptotic cell membranes. Cytotox integrates into cells with compromised membrane integrity and binds DNA.
3. Treat each well with 100 µL of 2X dye-dosed Organoid Culture Media.
4. Add 200 µL of sterile 1X PBS to the outer empty wells of the plate. Incubate at 37 °C overnight.
  1. *Note: PBS in the peripheral wells decreases evaporation of media from the inner wells*.
5. Addition of drugs/treatment agents (Day 0): Prepare drugs in pre-warmed Organoid Culture Media at a 2X concentration in a volume of 100 µl per well.
  1. *Note: DMSO can be toxic to cells at high concentrations. A concentration of 0*.*1% DMSO is not exceeded in our experiments. In addition to drugs, some fluorescent reagents are distributed as a DMSO solution. It is important to account for total DMSO concentration when working with such reagents*.
6. Add 100 µl of 2X treatment media to each well, avoid creating bubbles.

### Setting up imaging parameters

1. Launch Gen5 software to begin imaging the 96-well plate.
2. Place plate in Cytation 5. **Open** Gen5 software. **Click** New Task > Imager Manual Mode. **Select** Capture Now and input the following settings: Objective (select desired magnification); Filter (select microplate); Microplate format (select number of wells); and Vessel type (select plate type). **Click** “Use Lid” and “Use slower carrier speed.” **Click** OK.
  1. *Note for Vessel type: Be as specific as possible when selecting information about the plate because the software is calibrated to the specific distance from the objective to the bottom of the plate for each plate type and thickness of the plastic*.
  2. *Note for Slower Carrier Speed: Select this box to avoid disrupting domes when loading/unloading plates*.
3. Create a Z-Stack which will image the entire BME dome.
  1. **Select** a well of interest to view (left panel, below histogram).
  2. **Select** the Bright Field channel (left panel, top). **Click** Auto-expose and adjust settings as needed.
  3. Set the bottom and top of the Z-Stack: **Expand** Imaging Mode tab (left panel, middle). **Check** the Z-Stack box. Using the course adjustment arrows (left panel, middle), **click** the down adjustment until all PDOs have come into and then out of focus and are fuzzy. **Set** this as the bottom of the Z-Stack. Repeat in the opposite direction using the course adjustment arrows to set the top of the Z-Stack.
  4. To ensure that the Z-Stack settings are appropriate for other wells of interest, **select** another well (left panel, below histogram) and visualize the top and bottom of the Z-Stack. To manually enter the focal positions, **click** on the three dots next to the fine adjustment (left panel, top). A window will open; type in the top Z-Stack value (found in the left panel, center, under Imaging Mode). Repeat for the bottom Z-Stack value. Adjust as necessary to capture the desired Z range by repeating step 3. If adjustments were necessary, **select** another well to verify settings.
4. Set the exposure settings for fluorescent channel(s). Settings are described for two fluorescent channels (GFP & TRITC). The specific number of fluorescent channels will depend on the experiment and which fluorescent cubes are installed in the Cytation 5.
  1. *Note: If the signal intensity is anticipated to be significantly higher at the end of the experiment, users should consider performing test experiments to determine the optimal exposure settings at the end of the experiment that can then be applied when setting up the initial parameters*.
  2. **Expand** the Imaging Mode tab (left panel, middle) and open Edit Imaging Step. A pop-up window will appear.
  3. Under Channels, **click** on the bubble for the desired number of channels. One channel should be designated for Bright Field and additional channels for each fluorescent channel. In this example, Channel 1 = Bright Field; Channel 2 = GFP; Channel 3 = TRITC. Using the drop-down Color menus, **select** the appropriate setting for each channel. Close editing window by **clicking** OK.
  4. Set up each fluorescent channel.
    1. Switch the channel to GFP (left panel, top).
    2. **Click** Auto-expose (left panel, top). **Expand** the Exposure tab (left panel, middle) and adjust the exposure settings to minimize background signal.
    3. Copy exposure settings to the Image Mode tab.
      1. **Click** on the “Copy” icon 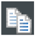 next to the Edit Imaging Step box.
      2. **Click** Edit Imaging Step, which will open another window.
      3. Under the GFP channel, **click** the “Clipboard” icon 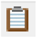 in the Exposure line. This function will add the Illumination, Integration Time and Camera Gain settings to the channel.
    4. Repeat Steps 1-3 for the TRITC channel.
    5. **Click** OK to close the window.
5. Set up the Image Preprocessing and Z Projection steps, which will automate image preprocessing.
  1. **Click** on the “Camera” icon (left panel, bottom corner). A new window will open.
  2. Under ADD PROCESSING STEP (left panel, bottom), **click** on Image Preprocessing. A new window will open.
  3. On the Bright Field tab, **deselect** “Apply image preprocessing.”
  4. For each Fluorescent channel tab, make sure “Apply image preprocessing” is selected. **Deselect** “Use same options as channel 1” and **click** OK. The window will close.
  5. Under ADD PROCESSING STEP, **click** on Z Projection. A new window will open. If desired, the slice range can be adjusted (e.g., to narrow the Z range). Close window by **selecting** OK.
6. Create Protocol.
  1. **Click** “Image Set” in the toolbar. In the drop down menu, **click** “Create experiment from this image set.” The imaging window will close and the Procedure window will open.
    1. *Note: The parameters selected in Imager Manual Mode will automatically be loaded into the new window whereby an experimental protocol can be created*.
  2. Set the temperature and gradient: **Click** on Set Temperature under the Actions heading (left). A new window will open. **Select** “Incubator On” and manually enter the desired temperature under “Temperature.” Next, under “Gradient,” manually enter “1.” Close window by **selecting** OK.
    1. *Note: Creating a 1 °C gradient will prevent condensation from forming on the lid of the plate*.
  3. Designate wells to image.
    1. Double **click** on the Image step under description.
    2. **Click** “Full Plate” (right corner, top). This will open the Plate Layout window.
    3. Highlight wells of interest using the cursor. **Click** OK.
    4. If desired, **check** “Autofocus binning” and “Capture binning” boxes. **Click** OK to close window.
      1. Note: Binning will require exposure adjustment, as described in Step 2.2 above. Please refer to *Data Management* in the Discussion for specific scenarios in which this feature may be used.
  4. Set intervals for kinetic imaging.
    1. **Click** on Options under the Other heading (left).
    2. **Check** the “Discontinuous Kinetic Procedure” box.
    3. Under Estimated total time, enter the run time for the experiment (e.g., 5 days). Under Estimated interval, enter the interval at which to image the plate (e.g., every 6 hours).
    4. **Click** “Pause after each run” to allow time for the plate to be transferred to the BioSpa incubator. Close window by **selecting** OK.
  5. Update Data Reduction steps.
    1. **Click** OK to close the Procedure window. A tab will open to update data reduction steps. **Select** Yes.
    2. Double **click** on Image Preprocessing. **Click** through the different channels to verify settings. **Click** OK.
    3. Double **click** on Z Projection. **Click** through the different channels to verify settings. **Click** OK.
    4. **Click** OK to close the Data Reduction window.
  6. Format the Plate Layout.
    1. Open the Plate Layout Wizard and designate well types.
      1. **Click** on the Plate Layout icon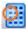 in the toolbar (left corner, top). This will open the Plate Layout Wizard.
      2. **Check** the boxes next to the well types used in the experiment. Under Assay Controls, enter the number of different control types using the arrows. **Click** Next. This will open the Assay Control #1 window.
    2. Set Assay Control well conditions.
      1. On the Assay Control #1 Window, enter control label in the Plate Layout ID box. If desired, enter the full name in adjacent box. **Select** the number of replicates for the respective control condition using the arrows.
      2. If using multiple concentrations or a dilution series within the control, **click** “Define dilutions/concentrations” and use the drop-down menu to **select** the Type. Enter values for each concentration/dilution in the table.
        1. *Note: The auto function can be used if concentrations change by a consistent increment*.
      3. **Select** the Color tab in toolbar. Choose desired Text color and Background Color for control in drop-down menu. **Click** Next.
      4. Repeat as necessary with additional controls.
    3. Set Sample well conditions.
      1. On the Sample set-up page, enter the sample ID Prefix (e.g., SPL). **Select** number of Replicates using the arrows. If using samples with varying treatment concentrations, **select** Concentrations or Dilutions in the Type drop-down menu. Enter dilutions/concentrations in the table and enter units in the Unit box.
      2. **Select** Identification Fields in the toolbar. Enter desired Category Name(s) (e.g., sample ID, drug) in the table.
      3. **Select** the Color tab in toolbar. **Select** a different color for each treatment group/sample.
        1. *Note: The numbers on the left side correlate with the different sample numbers*.
      4. **Click** Finish. This will open the Plate Layout page.
    4. Assign Sample IDs.
      1. **Select** SPL1 from the left panel. Use cursor to **select** wells.
        1. *Note: Autoselect tools can be adjusted in the serial assignment box. Number of replicates and orientation of layout can be pre-designated*.
      2. Repeat with other samples to complete plate layout. Once satisfied, **click** OK.
      3. In file toolbar, **select** Sample IDs. Fill in Sample ID columns with the appropriate information for each Sample (e.g., drug type). Press OK.
  7. Save the Protocol.
    1. In the toolbar, **Click** File > Save Protocol as.
    2. **Select** location to save file. Enter file name. **Click** Save to close window.
    3. **Click** File > Exit in the toolbar. A tab will open to save changes to Imager Manual Mode. **Select** No.
    4. A tab will open to save changes to Experiment 1. **Select** No.
    5. A tab will open to update the protocol definition. **Select** Update.
    6. **Close** Gen5 software.
7. Import the Protocol into BioSpa OnDemand and finish setting up the Experiment.
  1. **Open** the BioSpa OnDemand software.
  2. **Select** an available slot in the BioSpa.
  3. Remove the plate from the Cytation 5. **Click** Open drawer to access the appropriate slot in the BioSpa and insert plate. **Click** Close drawer.
    1. *Note: This step can be performed at any point once the Protocol has been created in Step 5*.*7 above. However, the plate must be in the Cytation 5 to perform a timing run in the below Step 6*.*4*.*5*.
  4. Import the Protocol.
    1. Under the Procedure Info tab, **select** User in the drop-down menu.
    2. Next to Protocol slot, **click** Select > Add a new entry.
    3. Next to Protocol slot, **click** Select. This will open a new window to navigate to the desired Protocol in the file architecture.
    4. **Click** Open to import the Protocol into the BioSpa OnDemand software.
    5. Enter the amount of time needed to image the plate. **Click** OK to close the Gen5 Protocol List window.
      1. *Note: This step is especially important when running several experiments at a time. To determine the time needed to image the*, ***click*** *“Perform a timing run now*.*”* ***Click*** *OK*.
  5. Set imaging intervals and schedule the experiment.
    1. Under Interval, enter the imaging interval which was designated previously in *Step 5*.*4* of *Setting up Imaging Parameters*.
    2. Under Start Time Options, **select** “When available.”
      1. *Note: A specific start time can be designated instead of running the protocol at the next available time*.
    3. Under Duration, **select** “Fixed” or “Continuous.”
      1. *Note: Selecting Fixed duration will set a specific endpoint for the experiment and requires the user to designate an experimental timeframe. Continuous duration will allow the experiment to run with no endpoint and can only be ended by a user stopping the experiment*.
    4. **Click** Schedule plate/vessel. This will open the Plate Validation Sequence. A tab will open with the proposed first read time. **Click** Yes to accept this schedule.

### Image analysis in Gen5 software (Figure 2)

1. Open image analysis module.

**Figure 2.**
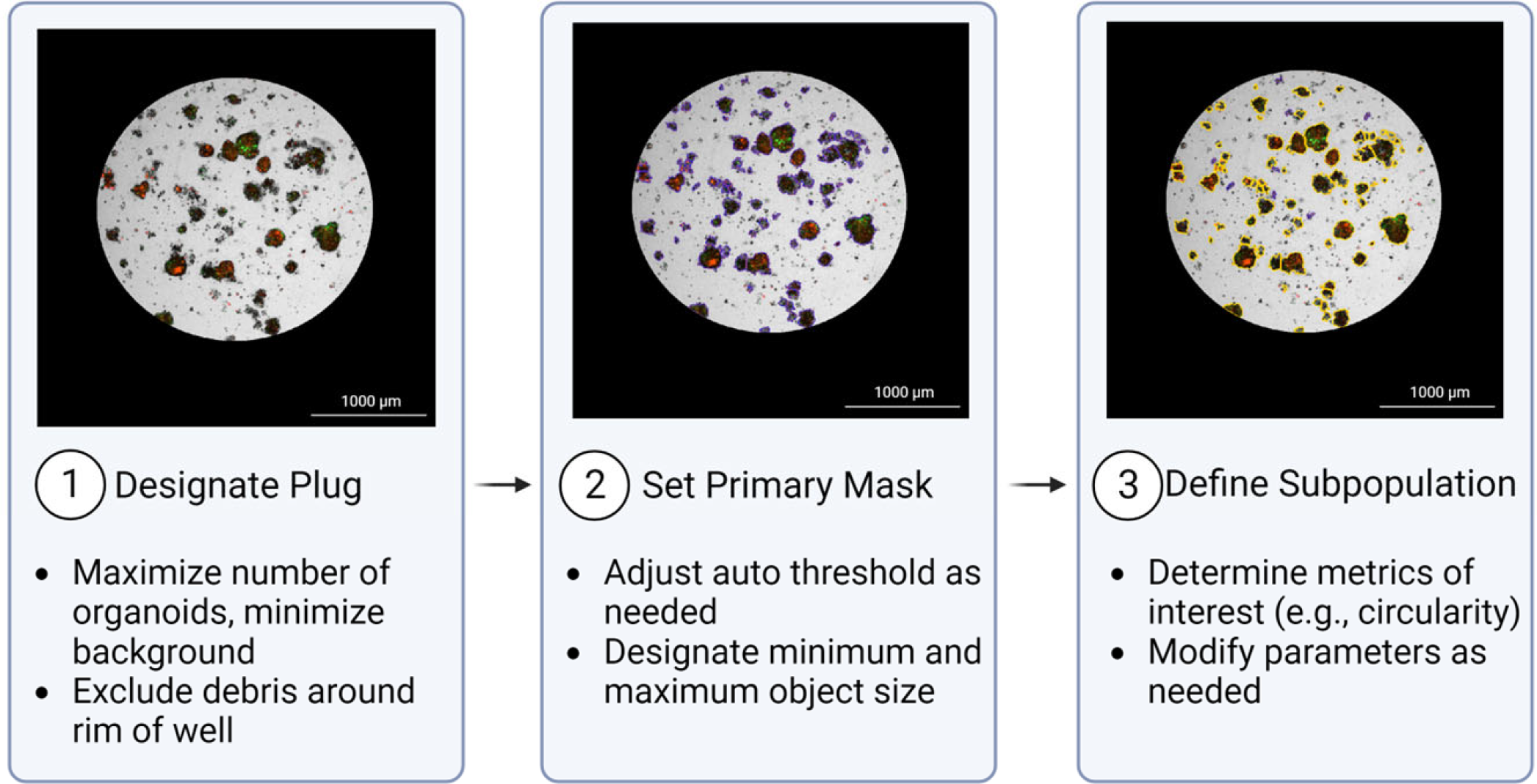
Overview of Cellular Analysis feature. 1: Designate Plug: A plug is designated to include areas of interest. 2: Set Primary Mask: The Primary Mask defines objects of interest based on size and pixel intensity in a channel of choice. In this representative image, objects included in the primary mask are outlined in purple. 3: Define Subpopulation: An additional subpopulation may be defined to further refine the desired population for analysis. The subpopulation in the example image (outlined in yellow) is defined based on circularity (>0.25) and area (>800). Images were acquired with a 4X objective.
  1. **Open** Gen5. In the Task Manager, **select** Experiments > Open. **Select** the experiment to open the file.
  2. **Click** Plate > View in the toolbar.
  3. Change Data drop-down menu to Z Projection.
  4. Double **click** on a well of interest.
  5. **Select** Analyze > “I want to setup a new Image Analysis data reduction step.” **Click** OK.
2. Cellular Analysis:
  1. Primary Mask
    1. Under ANALYSIS SETTINGS, **select** Type: Cellular Analysis and Detection Channel: ZProj[Tsf[Bright Field]] (left panel, center).
    2. **Click** Options. This will open the Primary Mask and Count page. In the Threshold box, **check** “Auto” and **click** Apply. **Click** the “Highlight Objects” box (right panel, bottom) to show objects within the designated threshold. Adjust as necessary to include objects of interest.
      1. *Note: Threshold settings are based on pixel intensity. For example, if the threshold is set to 5000, pixels with an intensity greater than 5000 will be included in the analysis*.
    3. Under Object selection, designate the minimum and maximum object size (µm). Adjust as necessary to exclude cellular debris/single cells.
      1. *Note: PDO size may vary significantly between different models and types. Use the measuring tool* 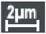 *in the Gen5 software to determine the minimum and maximum PDO size thresholds for each model. Users may choose a smaller minimum PDO size threshold relative to the value provided by the measuring tool in order to prevent exclusion of PDO fragments at later timepoints after drug treatment*.
    4. To limit the analysis to a certain region of the well, **deselect** “Analyze entire image” and **click** “Plug.” In the Image Plug Window, use the drop-down menu to select Plug shape. Adjust the size and position parameters as necessary to fit over the region of interest.
      1. *Note: It is important to maximize the number of PDOs within the plug while also excluding PDO-free areas to minimize background. Designate a plug size that will consistently capture the majority of the objects of interest across replicates. Generating a plug that also excludes the edges of the dome is important as it excludes any objects that may appear distorted due to the refraction of light from the extreme curvature of the dome around the edges*.
      2. *Note: “Include primary edge objects” may also be* ***deselected*** *to only capture entire PDOs within the plug*.
  2. Subpopulation Analysis. An example of subpopulation designation is provided in **Figure 3**.

**Figure 3.**
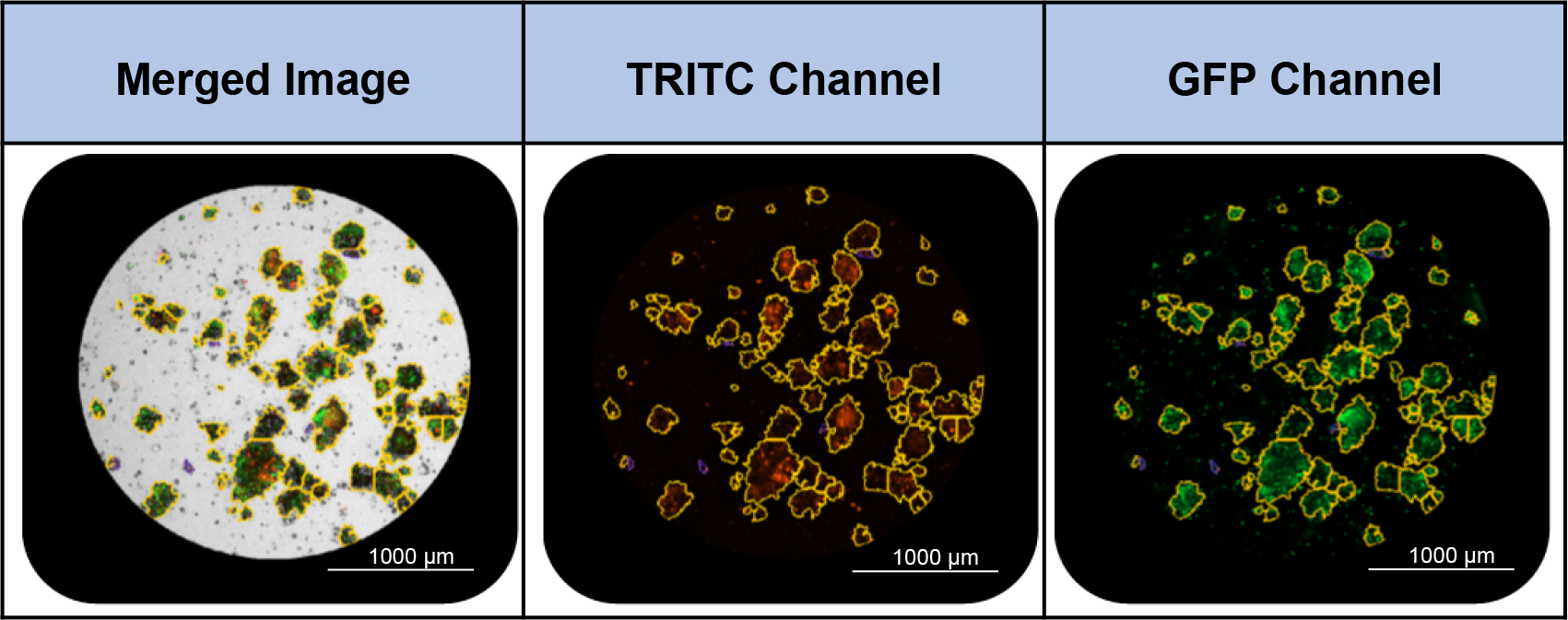
Examples of subpopulation masking using the Cellular Analysis feature. Subpopulations are defined in the Bright Field channel. The subpopulation in the example images (outlined in yellow) is defined based on circularity (>0.25) and area (>800). Images were acquired with a 4X objective.
    1. **Click** on Calculated Metrics in the Cellular Analysis toolbar. **Click** “Select or create object level metrics of interest” (right corner, bottom). Under Available object metrics, **select** metrics of interest (e.g., Circularity) and **click** the Insert button. **Click** OK.
      1. *Note: Morphology and density of each PDO model will determine the best metrics of interest to distinguish the subpopulation*.
    2. **Click** on Subpopulation Analysis in the Cellular Analysis toolbar. **Click** Add to create a new subpopulation. A pop-up window will open.
    3. If desired, enter a name for the subpopulation. Under Object metrics, **select** a metric of interest and press Add Condition. In the Edit Condition window, enter parameters for the chosen Object metric. Repeat with additional metrics as necessary.
      1. *Note: Parameters may be adjusted manually or set using the finder tool. For example, to exclude debris, users could add Area as an Object metric and select objects smaller than 800. We also routinely use Circularity as an Object metric and include any objects with a circularity greater than 0*.*2-0*.*5, depending on the model*.
    4. In the table at the bottom of the window, **check** the desired results to display. **Click** OK > Apply.
    5. To view the objects within the subpopulation, use the Object details drop-down menu (right panel, center) to select the subpopulation. Objects that fall within the parameters will be highlighted in the image.
      1. *Note: To change the highlight colors of the primary mask and subpopulation*, ***click*** *“Settings” (right panel, bottom)*.
    6. To adjust subpopulation parameters, reopen the Subpopulation Analysis window from the Cellular Analysis toolbar. **Select** the subpopulation and **click** Edit.
    7. **Click** ADD STEP.
      1. *Note: This will apply the same analysis to all wells within the experiment at all time points. In the drop-down menu on the Matrix page, different metrics can be selected for individual viewing*.

### Exporting data from Gen5 to Excel

1. To customize a data file for export, **select** the Report/Export Builders icon 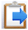 in the toolbar. In the pop-up window, **click** New export to Excel.
2. On the Properties page of the pop-up window, **select** Scope > Plate and Content > Custom. **Click** on the Content option in the toolbar. **Click** Edit template, which will open the Excel program.
3. Within the Excel program, **select** Add-ins > Table > Well Data. Hover over the various selections to see options for export. **Select** metric of interest (e.g., Object Mean[ZProj[Tsf[TRITC]]]).
  1. *Note: Plate layout can be added to the Excel analysis template by selecting Add-ins > Protocol Summary > Layout*.
4. An Edit window will open. In the Wells box, designate the wells for export either by Well-ID or Well #. **Select** OK. A template will be loaded into the Excel file. **Close** Excel program. The template is automatically saved.
5. **Click** OK on the New export to Excel window and **Close** the Report/Export Builders window.
6. **Click** the Export icon 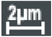 in the Gen5 toolbar. **Check** the box next to the desired Export file. **Click** OK. Gen5 will automatically populate the Excel template and open the file in Excel.

### External data analysis

1. **Open** the Export file (.xlsx) in Excel.
2. For each well, divide each individual value by the 0:00 time point value for that well. This will set time point 0 equal to 1 and each value beyond that will be relative to the initial reading.
3. **Open** a new file in the GraphPad Prism software. In this protocol, version 9.5.1 was used. **Select** the XY layout option.
4. Input labels for each data group. Copy and paste the time points and corresponding normalized values for each treatment group into the Prism table. A graph for the data will be automatically generated and can be found under “Graphs.”

## REPRESENTATIVE RESULTS

Our objective was to demonstrate the feasibility of using multiplexed live-cell imaging to assess PDO therapeutic response. Proof of concept experiments were performed in two separate PDO models of endometrial cancer: ONC-10817 and ONC-10811 (see **Supplementary Figure S1 & S2** for ONC-10811 data). Apoptosis (annexin V staining) and cytotoxicity (Cytotox Green uptake) were kinetically monitored in response to the apoptosis-inducing agent, staurosporine. Specifically, PDOs were plated in 96-well plates, treated with Annexin V Red and Cytotox Green dyes, and placed in a 37 °C incubator overnight as diagrammed in **Figure 1**. We confirmed in two independent PDO models that treatment with Annexin V and Cytotox Green dyes is not toxic (**Supplemental Figure S2**). The following day, PDOs were treated with increasing concentrations of staurosporine (0.01 nM, 0.1 nM, 1 nM, 10 nM, 100 nM, 500 nM). Subsequently, protocols were established in the Gen5 software and experiments were set to run over a period 5 days, imaging every 6 hours. Data were analyzed using the Cellular Analysis function in the Gen5 software as described in the *Image Analysis in Gen5 software* protocol. The Primary Mask was set using the Auto threshold function with “Split touching objects” unchecked and with size parameters of minimum: 30 µm and maximum: 1000 µm. The PDO subpopulation was defined by circularity > 0.25. The Object Mean Intensities in the TRITC (annexin V, apoptosis) and GFP channels (Cytotox Green, cytotoxicity) within the designated PDO subpopulation were exported as an .xlsx file for further analysis. The Object Mean Intensity for each well at each time point in both the GFP and TRITC channels was normalized to time 0. Normalized fluorescence data were then transferred to a Prism file and visualized as a line plot.

Treatment with staurosporine resulted in a significant, dose-dependent increase in apoptosis and a decrease in cell health over time in comparison to vehicle control, as evidenced by the increase in Object Mean Intensity in both the TRITC (annexin V) and GFP (Cytotox) channels (**Figure 4, Figure 5** and **Supplemental Videos SV1-7**). The 500 nM, 100 nM, and 10 nM doses of staurosporine each resulted in a statistically significant increase in both apoptosis and cytotoxicity over time (**Figure 5A-C**) as well as at the end of the experiment (**Figure 5E**,**F**). Furthermore, staurosporine effectively inhibited PDO growth and formation at these concentrations as demonstrated by an overall decrease in total PDO area, whereas the control wells exhibited an increase in PDO area (**Figures 4 & 5D**).

**Figure 4.**
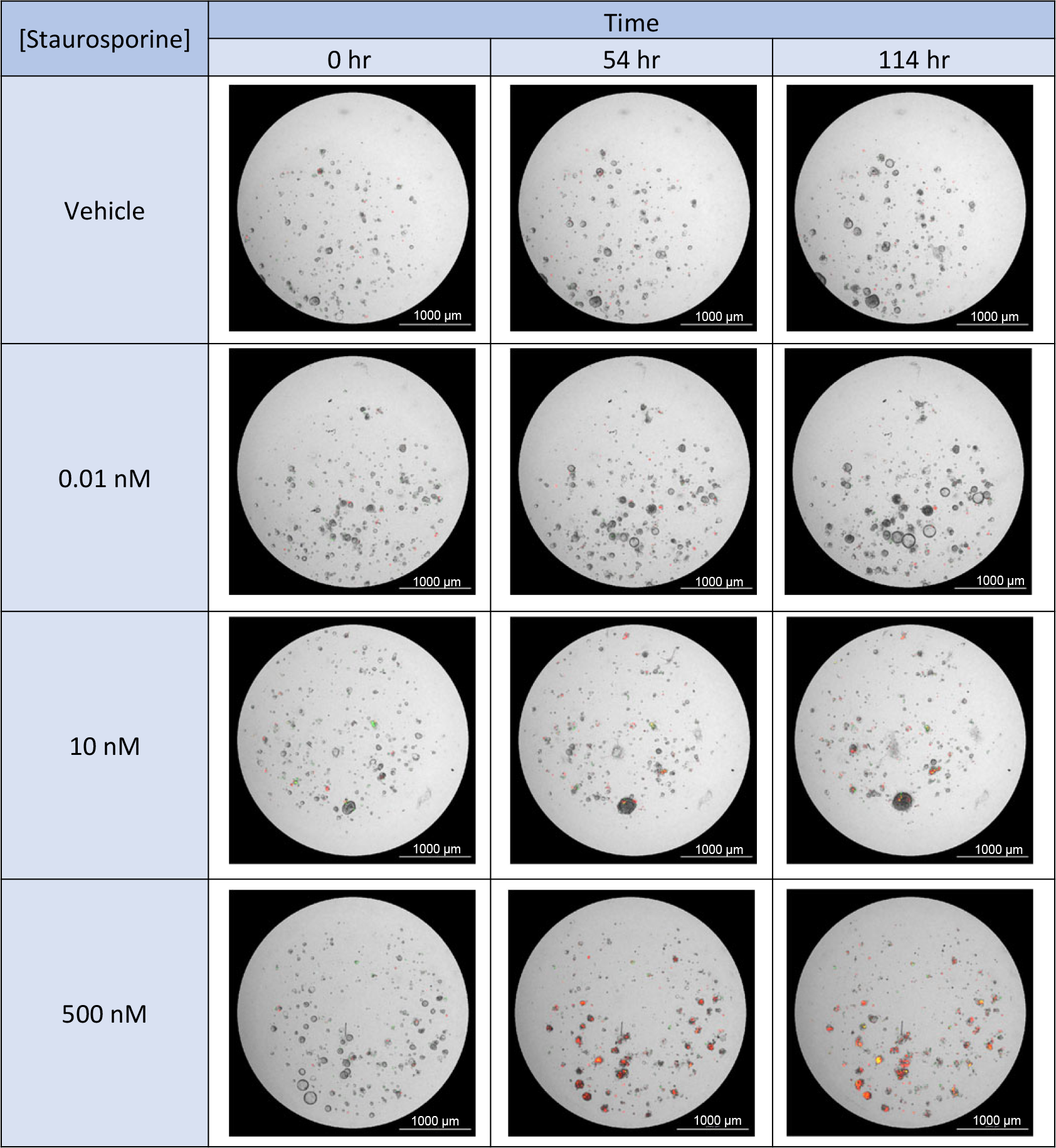
Staurosporine treatment results in a dose‐dependent increase in apoptosis and cytotoxicity. Bright Field images (4X objective) with GFP and TRITC fluorescence overlay are shown for **the 500 nM, 10 nM, and 0.1 nM doses of staurosporine at 3 time points**: 0 hr, 54 hr, 114 hr. Red fluorescent signal indicates apoptosis (annexin V) and green fluorescent signal indicates cytotoxicity (Cytotox).

**Figure 5.**
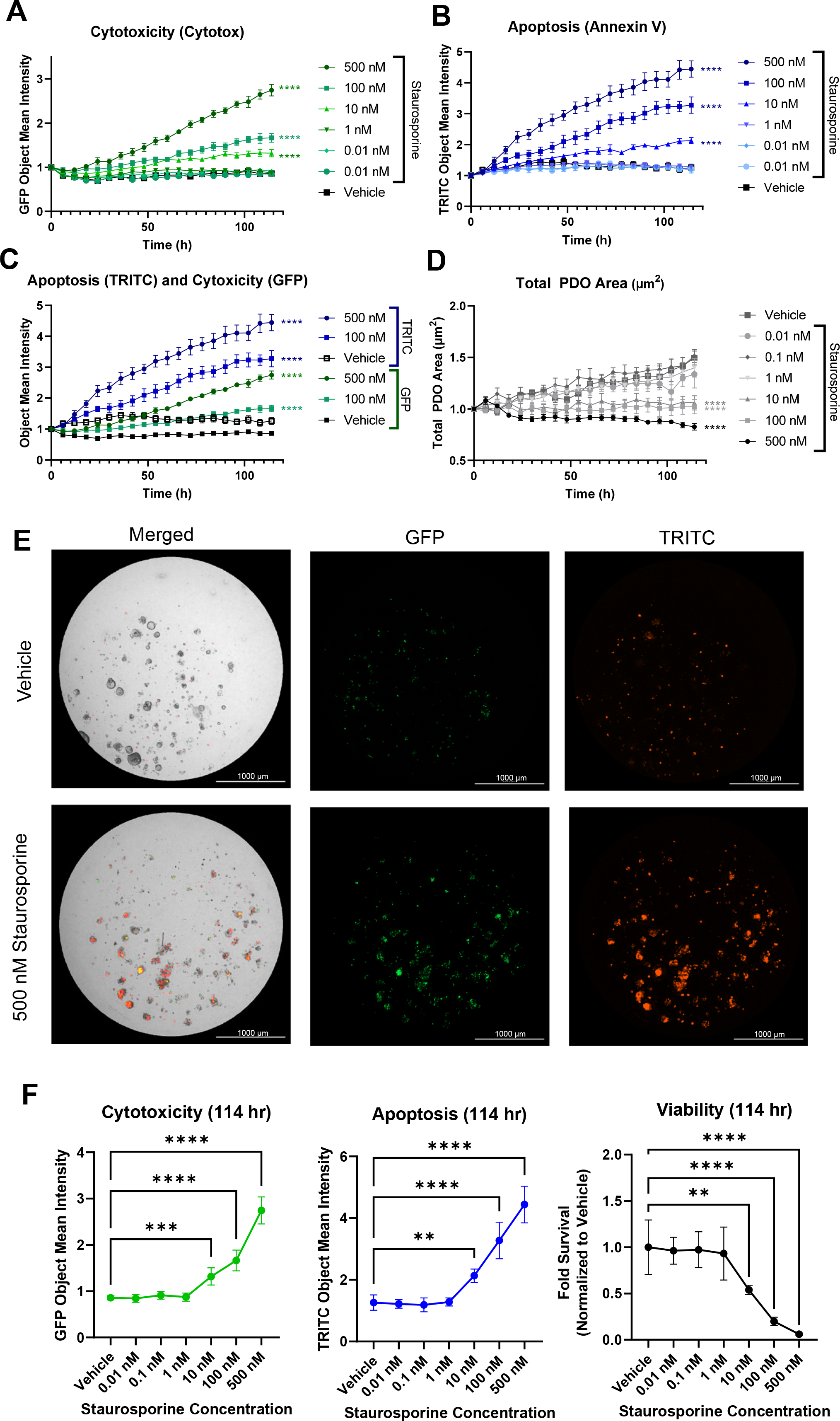
Multiplexed fluorescent live‐cell imaging to assess PDO response. PDO model ONC-10817 was plated in 96-well plates and incubated with Annexin V Red (1:400) and Cytotox Green (200 nM) dyes overnight at 37 °C. The following day, PDOs were treated with increasing concentrations of staurosporine and were imaged every 6 hours for ∼5 days. (**A, B**) Time and dose-dependent increase in cytotoxicity (**A**) or apoptosis (**B**) in response to staurosporine. Data were plotted as the Object Mean Intensity in the GFP or TRITC channel. **(C)** Comparison of time course of apoptosis and cytotoxicity in response to 100 nM or 500 nM staurosporine. Data were plotted as the Object Mean Intensity values in the GFP and TRITC channels. (**D**) Staurosporine inhibits growth of PDOs. Data were plotted as the average total PDO area. Data in A-D were normalized to PDO number at time 0 in each well and plotted as the mean and standard error of the mean (SEM). N=5 technical replicates per treatment. **** p < 0.0001 vs. vehicle control by 2-way ANOVA. (**E**) Representative Bright Field, GFP, and TRITC images of 500 nM staurosporine-treated PDOs vs. vehicle at the end of the experiment (114 hr). Images were acquired with a 4X objective. (**F**) Quantification of cytotoxicity, apoptosis, and viability at the 114 hr timepoint. GFP Object Mean Intensity (left) and TRITC Object Mean Intensity (middle) were calculated at the 114 hr timepoint using results from panels A-C. Viability (right) was assessed using the CellTiter-Glo 3D reagent per the manufacturer’s protocol. Raw luminescence (RLU) values were normalized to total PDO area at time 0 and plotted as the fold viability relative to vehicle control, which was set at 1.0. ** p<0.01, ***p<0.001, **** p<0.0001 vs. vehicle control via one-way ANOVA with Dunnett’s post-hoc test. N=5 technical replicates per treatment.

Since a major advantage of live-cell imaging is the ability to correct for variability in plating, we performed an experiment whereby cell viability was assessed as an endpoint measure. Proof-of-concept endpoint assay data were collected using a PDO model that was generated from a patient-derived xenograft of prostate cancer. Bright Field images were collected at the beginning of the treatment period (day 0), followed by addition of a dual dye reagent that measures both viability (acridine orange, AO) and cell death (propidium iodide, PI). The AO component emits a green fluorescent signal upon binding to double stranded DNA, an indicator of cell viability. The PI component stains dead nucleated cells and can be used to quantitate cell death in response to treatment. In order to account for variability in PDO plating, we devised a method to determine the number of PDOs per well at time 0 by converting Bright Field images to Digital Phase Contrast images (**Supplemental Figure S3** and **Supplemental Protocols**).

Prostate cancer PDOs were treated with daunorubicin, a chemotherapy agent that causes cell death, for 7 days. Upon completion of the experiment, samples were stained with AOPI as described in **Supplemental Protocols**, followed by analysis of fluorescent images in Gen5. **Figure 6A** shows a panel of images from the AOPI endpoint assay on day 7. When comparing vehicle-treated PDOs (row one) to PDOs treated with 10 µM daunorubicin (row two), there was a clear decrease in green fluorescence (measure of viability, column two) and an increase in red fluorescence (measure of cell death, column three). These results were then quantitated in **Figure 6B**, where we show the array of readouts that can be achieved using the AOPI endpoint staining technique. The upper left plot depicts the viability measurement generated from the AO stain, normalized to the PDO count determined by Digital Phase Contrast image analysis of each well on Day 0. These data correlate with the visual result from **Figure 6A**, whereby as the concentration of daunorubicin increased, the viability drastically decreased. This is further recapitulated in the upper right graph which demonstrates an increase in cell death denoted by an increase in red fluorescence acquired with the PI stain.

**Figure 6.**
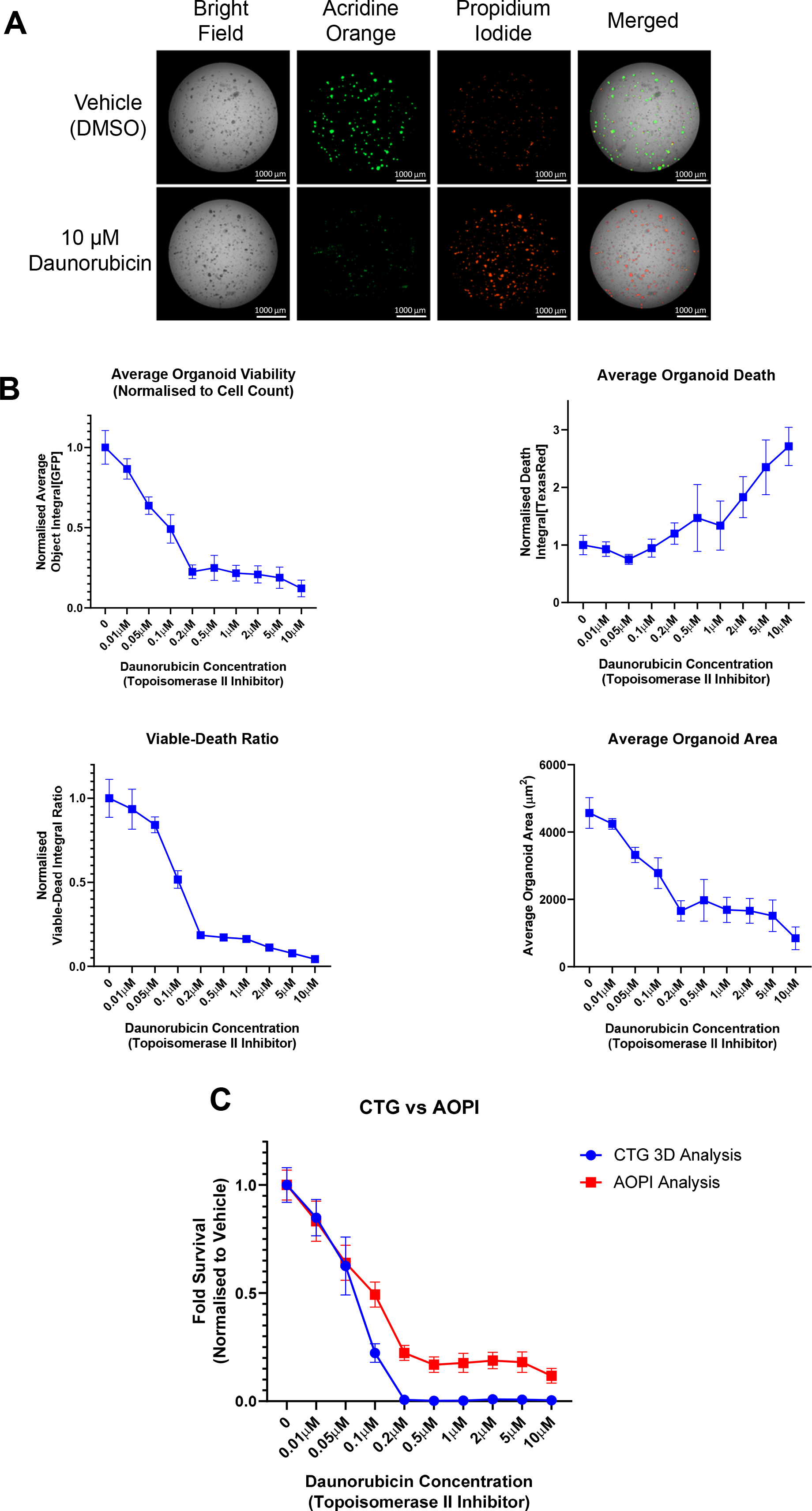
Use of live‐cell imaging to aid in normalization of endpoint assay data. PDOs were treated with increasing concentrations of the topoisomerase-II inhibitor, daunorubicin, for 7 days. PDOs were exposed to AOPI Staining Solution and imaged as described in **Supplemental Protocols**. AO= acridine orange, a measure of viability (GFP channel); PI= propidium iodide, a measure of cell death (Texas Red channel). (**A**) Representative images acquired using AOPI staining after 7 days of treatment with 10 µM daunorubicin or vehicle control (0.1% DMSO). (**B**) Different readouts using AOPI fluorescence as an endpoint viability/cell death method. See **Supplemental Protocols** for a detailed description of the analysis methods. Upper left, analysis of PDO viability after 7 days as determined by AO staining. Upper right, analysis of cell death by PI staining. Data for AO and PI staining were normalized to PDO number at time 0 and then to vehicle control, which was set at 1.0, and plotted as the mean and standard deviation. Lower left, calculation of viable to dead ratio using the average object integrals for the AO and PI stains. Lower right, area of PDOs as determined by AO staining. Cellular Analysis was performed in the GFP channel. (**C**) Comparison of two methods to test PDO viability. After imaging, viability was evaluated using the CellTiter-Glo 3D reagent per the manufacturer’s protocol. The fold survival relative to vehicle control was plotted at increasing concentrations of daunorubicin. Data represent the mean and standard deviation for N=6 technical replicates per treatment; data were not normalized to time 0 in panel C.

The PI data were then combined with the viability reading (AO) to calculate a Viable to Dead Ratio (**Figure 6B**, lower left graph). This ratio is a useful approach to determine whether a drug is cytostatic or cytotoxic. Specifically, a cytotoxic drug will reach much closer to 0 than a cytostatic drug, due to the fact a drug that is cytostatic will inhibit growth but may not induce cell death. Lastly, the area of the PDOs can be accurately calculated using the green fluorescence of the AO stain, even when PDOs may be undergoing cell death and blebbing. The lower right graph depicts the average PDO area, which was calculated as the sum of the area denoted in the subpopulation analysis divided by the PDO number. Analysis of area can give further indication as to whether a treatment is simply inhibiting PDO growth or actually causing PDO regression. Note that the analysis of average PDO area was performed using the GFP channel and Cellular Analysis function, in contrast to **Figure 5D** that used the Bright Field images to calculate total PDO area. These data highlight the flexibility of the analysis pipeline depending on data availability and user interest.

Finally, we compared the gold standard for viability readings, CellTiter-Glo 3D, to the viability fluorescence reading using AOPI (**Figure 6C**). Note that the data in this panel were not normalized to the time 0 PDO number since this normalization is not typically performed by labs using the CellTiter-Glo 3D kit. We observed the same trend for drug effect in both assays, whereby PDO viability decreased as the daunorubicin concentration increased. The only visual difference between these readouts was that the CellTiter-Glo 3D analysis reached an IC50 before the AOPI analysis and nearly completely reached 0. This result may be explained by the mechanism of action of daunorubicin. Daunorubicin is a topoisomerase-II inhibitor that introduces double stranded DNA breaks, leading to cell cycle arrest and eventually apoptosis^14^. During cell cycle arrest, ATP depletion can occur^15^. Given that the CellTiter-Glo 3D assay is based on an ATP-luciferase reaction to generate a luminescence signal, we hypothesize that the stronger reduction in cell viability at higher concentrations of daunorubicin was due to ATP depletion rather than complete cell death. Supporting this idea, the images in **Figure 6A** depict a population living PDOs in the culture, as denoted by green fluorescence.

## DISCUSSION

PDO cultures are becoming an increasingly popular in vitro model system due to their ability to reflect cellular responses and behaviors^2^. Significant advances have been made in PDO generation, culture and expansion techniques, yet methods to analyze therapeutic responses have lagged. Commercially available 3D viability kits are lytic endpoint assays, missing out on potentially valuable kinetic response data and limiting subsequent analyses by other methods^8^. Emerging studies are applying live-cell imaging to assess drug responses in PDO models. Here, we present a method to assess PDO therapeutic responses over time utilizing multiplexed fluorescent live-cell imaging. Multiplexing fluorescent dyes allows for different cellular responses to be quantified simultaneously. In addition to apoptosis and cytotoxicity, we envision that this approach can be expanded in future studies to examine other phenotypic effects in PDOs.

### Critical steps within the protocol

The protocol presented herein is designed to acquire kinetic bright-field and fluorescent images of PDOs plated as domes in a 96-well plate. Key steps include 1) plating; 2) treating with dye and drug; 3) defining imaging parameters; 4) image preprocessing and analysis at the completion of kinetic image acquisition; and 5) data export for analysis in other statistical software (**Figure 1**). Intact PDOs are plated in a 96-well plate as 5 µL BME domes composed of a predefined ratio of BME and Organoid Culture Media (typically 1:1). The protocol presented herein uses UltiMatrix as the BME due to minimal batch-to-batch variability and superior optical properties for imaging. Modifications for harvesting PDOs cultured in other BMEs such as Matrigel may be necessary, as well as for models that are clumpy and difficult to dissociate (see examples **Supplemental Figure S4**). Next, fluorescent dyes are added to the Organoid Culture Media at the time of plating (Day -1) at a 2X concentration in a 100 µl volume and incubated overnight to determine baseline cell death. The following day (Day 0), drugs or other treatments are added to Organoid Culture Media at a 2X concentration in a 100 µl volume, and then added to each well for a final volume of 200 µl. The final volume of 200 µl reduces the meniscus effect with imaging. Kinetic images are acquired using the Cytation 5 plate reader partnered with the BioSpa tabletop incubator. The BioSpa incubator allows for incubation of experimental plates at a fixed temperature of 37 °C and 5% CO_2_ environment. Up to 8 plates can be stored in the BioSpa and thereby 8 experiments can be simultaneously conducted. Imaging parameters for each plate are set up in the Gen5 software using “Imager Manual Mode” and saved as a Protocol. The Protocol is then loaded into the scheduling software (BioSpa OnDemand). The BioSpa OnDemand software automates the workflow for kinetic image acquisition, including physical transfer of the plate from the BioSpa incubator into the Cytation 5 and dictating which protocol to run in Gen5 for data collection. Images are analyzed using the Cellular Analysis function in the Gen5 software (**Figure 2**). We describe specific methods to analyze Z Projections of fluorescent and Bright Field images. Specific methods to analyze single Z planes and/or only Bright Field images are provided in **Supplemental Protocols**. Finally, users can define specific data features, such as PDO area, number, and mean fluorescence intensity, to export as an Excel spreadsheet for subsequent statistical analysis of drug effects.

### Significance with respect to existing methods

Given that PDO models are a relatively new model system to interrogate drug effects, methods to accommodate the culture conditions, in particular growth in BME, are still emerging^16^. The bulk of studies assessing PDO response to drug treatment rely on ATP-based endpoint assays as a surrogate for cell health (CellTiter-Glo 3D). This method requires cell lysis, thus precluding subsequent downstream analyses. Alternative endpoint assays, such as fluorescent staining, single timepoint imaging, and morphologic tracking, have provided other metrics to characterize drug response while allowing for sample use for additional purposes. For example, in-plate endpoint fixation protocols have been applied for high-throughput analysis of drug effects^17,18.^ An advantage of this method is that it circumvents the extensive processing required in typical immunohistochemistry and immunofluorescent imaging^19^ and is particularly useful when sample is limited, as is the case for PDO models. It is also amenable to high-resolution imaging with confocal microscopy. Another endpoint assay that does not require cell lysis is live-cell imaging with reagents such as propidium iodide or acridine orange. Our comparative analysis of CellTiter-Glo 3D and AOPI, the latter of which does not require cell lysis, for assessment of cell viability and death (**Figure 6C**) highlights the advantages of using live-cell imaging dyes in PDO models. This method has been applied by several groups, including ours, to assess phenotypic effects at the conclusion of an experiment^18-20^. However, kinetic data acquisition using either the in-well fixation or endpoint live-cell imaging methods require significant sample. Label-free morphologic assessment of PDOs over time in part overcomes this limitation and can be assessed using a wide range of imaging modalities^8,10^,17,18, yet changes in morphology may not be representative of the spectrum of potential drug effects such as apoptosis and changes in cell viability. We have observed significant model-to-model variability in morphologic changes in response to drugs. For example, some PDOs will increase in area as they are undergoing apoptosis, whereas other models may shrink. Kinetic morphologic assessments have been multiplexed with either fixed or live-cell fluorescent endpoint assays in a limited number of studies^18,20.^The methods presented herein are among the first to multiplex different fluorescent dyes to simultaneously assess multiple cellular effects in a high-throughput system.

### Method troubleshooting and modifications

One of the greatest improvements provided by kinetic live-cell imaging is that it overcomes key limitations associated with endpoint assays. For example, plating an equal number of organoids per well is technically difficult because most automated cell counters are gated for objects smaller than 60 µm. Live-cell imaging allows for normalization to PDO number or area at time 0 in each well, which can then be used to adjust for variation in plating amongst wells. The gold standard endpoint assay for PDOs is the CellTiter-Glo 3D kit, which measures ATP as a surrogate for cell viability. We have extensively used this assay in our studies of gynecologic and prostate cancer PDOs^13,21-24.^ In addition to requiring cell lysis for ATP measures, drugs and other therapeutic modalities can alter ATP levels, potentially providing an inaccurate assessment of therapeutic response. Utilizing live-cell imaging dyes allows for quantification of a wider scope of specific cellular responses, such as apoptosis and cytotoxicity. Importantly, these reagents do not perturb cell viability or induce DNA damage, as is the case with propidium iodide; this allows for repeated assessment of PDO response over time. While we have not explored the use of acridine orange (AO) for kinetic measures, the mechanism of action for AO should not preclude such application in the future.

Tumors consist of several distinct cell populations with varying genomic profiles and morphologies. Due to the complex heterogeneity of PDO models, individual organoid responses may vary and tracking these responses is challenging. Our protocol provides a method to assess multiple organoid response metrics on a well-by-well basis. However, certain technical considerations still apply for live-cell imaging. The number of wells of a 24-well plate needed to seed a 96-well plate will depend upon the density and growth rate of each PDO model. Because clumping can confound image analysis (**Supplemental Figure S4**), PDO models may need additional processing steps prior to plating. If necessary, PDOs may be mechanically sheared during processing through vigorous pipetting with a non-wide bore p200 tip. Enzymatic digestion using TrypLE Express may also be employed prior to plating to promote PDO dissociation and decreased clumping. In our experience, we have noted that the length of TrypLE incubation is highly variable depending upon the model. Therefore, TrypLE incubation should be optimized for each model, particularly if researchers wish to obtain single cell suspensions. Recovery time prior to beginning treatment may be necessary for PDO models that are particularly sensitive to enzymatic digestion.

We present methods for specific live-cell imaging reagents. A limitation in the broad applicability of the methods presented herein is a paucity of reagents that are compatible with BME. For other dyes/reagents that have not yet been optimized for use in PDOs, additional troubleshooting may be necessary to determine the optimal concentration and minimize solvent effects. Depending on the readout of interest (e.g., apoptosis or cytotoxicity), treatment with a compound known to induce cell death, such as staurosporine or daunorubicin^13^ should be used to optimize reagent conditions. An additional consideration is the optimal time for dye integration into the BME dome. In our experiments, we perform an overnight incubation prior to treatment to allow for complete uptake of the dye into the domes as well as determine baseline cell health. Since signal intensity will increase over time, researchers should perform pilot studies with the dyes to determine the ideal exposure time at the end of the experiment to avoid overexposed images. Finally, reagents should not interfere with cellular processes. For example, dyes that intercalate into DNA are not compatible with kinetic imaging. However, live-cell imaging can be used for data normalization in endpoint assays. Indeed, we present a supplementary method to quantify cell viability and the viable:dead cell ratio using AOPI. In this experiment, the fluorescent signal is normalized to PDO number for each well as determine by live-cell imaging on day 0 (day of treatment, **Figure 6**).

Another limitation in this methods paper is the reliance on manual pipetting for experiments, which can lead to greater variance in technical replicates. Further improvements in data reproducibility can be achieved with addition of automation to other steps of the experimental process, such as use of automated liquid handler for plating and drug dispensing as has been demonstrated by others^8,10.^However, this addition requires an additional investment in research infrastructure that may not be available to all investigators. Given the heterogeneity in PDO area for each model, normalizing results to the initial PDO number or area at time 0 is still a useful approach with automated seeding.

### Choosing metrics for analysis

A key advantage of the Cytation 5 over other live-cell imaging platforms is the ability to customize image and analysis features. However, the Cytation 5/Gen5 software has a steep learning curve and assumes that users have a foundational knowledge of imaging. One of our goals of this methods paper is to provide step-by-step instructions to decrease the barrier for other researchers to incorporate sophisticated live-cell imaging techniques in their PDO research programs. While the specific analysis steps presented herein are for one system, users can apply the multiplexing principles to other platforms, with the understanding that downstream analysis may necessitate use of third-party software, such as NIH ImageJ.

Analysis metrics should be chosen based on both experimental goals and plating conditions. For example, if the experiment is conducted using a fluorescent dye to quantify cell death, fluorescence intensity is an effective read-out. The most effective fluorescence metric (Total Fluorescence vs. Object Mean Intensity) will depend on the plating conditions (**Supplemental Figure S4**). If PDOs are evenly dispersed and are of a more circular, cohesive morphology, fluorescence can be determined in a defined PDO subpopulation. The specific function in the Gen5 software is Cellular Analysis and the output is Object Mean Intensity. However, if the PDO model is clumpy or has a discohesive morphology, we recommend quantifying the fluorescence signal at the image level; this function can be found under Image Statistics and the output is Total Fluorescence Intensity. While this method is useful for observing changes at the individual well level, this metric is not specific for objects of interest and could change erroneously if the PDOs move outside of the designated plug over the duration of the experiment. In scenarios in which there is significant debris or dead single cells, use of the Cellular Analysis is suggested to gate out any fluorescent signal that is not specifically associated with a PDO. An example of differential results using Cellular Analysis vs. Image Statistics is presented in **Supplemental Figure S5**.

To determine the optimal threshold for the Image Statistics function, the line tool may be utilized to determine the starting point. Using the line tool, users can determine the range of fluorescence intensity within an image. We set the background as the 25% value of the peak intensity within the image and designate this value as the lower threshold. To view the fluorescent areas that are included in the designated threshold, check the “Threshold Outliers” box. Additional fine-tuning of the lower threshold value may be necessary.

### Data management

A significant challenge in live-cell imaging is storing the massive amount of data generated with each experiment. This is particularly relevant in the case of PDO cultures, where Z-Stacks are used to image across several focal planes and generate a Z Projection. There are multiple methods to overcome this issue. First, ensure adequate storage capacity. We have installed three solid state hard drives with a total of 17 TB. Other options include transferring experimental files to external hard drives or networked storage. It is not recommended to directly write files to cloud based storage. To analyze data that has been stored on an external drive, simply transfer the experiment and the image files to a computer equipped with Gen5 software (NOTE: large files may take extended periods of time to transfer). Before analysis, the images must be re-linked to the experiment. Open the experiment file, click Protocol in the toolbar, select Protocol Options. Click Image Save Options and click Select new image folder. Locate the image file and click Select Folder to relink.

Depending on the goals of the experiment, it may also be suitable to use the binning feature within Gen5. Binning decreases file size by lowering the number of pixels, which leads to lower image resolution (see Step 5.3.4 in *Setting up imaging parameters* section). Therefore, binning is not recommended if high resolution images are required. When using the binning feature, the exposure settings will need to be adjusted. Once the experimental file has been created, double click on the Image section, and click the microscope icon to reopen Imager Manual Mode. Use the Auto-expose function or manually adjust exposure as needed.

## Conclusions

In summary, we present methods for assessing apoptosis and cell health of PDOs in response to cytotoxic agents. Future studies are necessary to optimize methods and develop additional analysis strategies for kinetic imaging of PDOs, such as other phenotypes and effects of drugs that are cytostatic rather than cytotoxic. A major roadblock is the commercial availability of dyes and reagents that are compatible with BME. There is still more work necessary to better understand how kinetic live-cell imaging can be fully utilized to extract the most data from these models.

## ACKNOWLEDGMENTS

We are grateful to the Tissue Procurement Core and Dr. Kristen Coleman at the University of Iowa for providing patient tumor specimens and to Dr. Sofia Gabrilovich in the Department of Obstetrics and Gynecology for assisting with patient-derived organoid model generation. We also thank Dr. Valerie Salvatico (Agilent, USA) for critical analysis of the manuscript. We acknowledge the following funding sources: NIH/NCI CA263783 and DOD CDMRP CA220729P1 to KWT; Cancer Research UK, Prostate Cancer UK, Prostate Cancer Foundation and Medical Research Council to JSdB. The funders had no role in the design or analysis of experiments or decision to publish.

## CONTENTS

### Supplemental Protocols

1. End of treatment cell viability and cell death fluorescence imaging using AOPI Staining Solution
2. End of treatment cell viability and cell death fluorescence image analysis in Gen5 software
3. Setting up imaging parameters for a single focal plane of view analysis (Bright Field/Digital Phase Contrast Images)
4. Digital Phase Contrast Image analysis in Gen5 software

### Supplemental Table

1. Table S1: Organoid Culture Media Components

### Supplemental Figures

1. Figure S1: Multiplexed live-cell imaging of ONC-10811.
2. Figure S2: Treatment with Annexin V and Cytotox does not perturb PDO viability.
3. Figure S3: Label-free analysis of PDOs using digital phase contrast.
4. Figure S4: PDO models may vary in their morphology and plating consistency.
5. Figure S4: Example images using Cellular Analysis vs. Image Statistics for quantifying fluorescence.

## SUPPLEMENTAL PROTOCOLS

**Setting up imaging parameters for a single focal plane of view analysis (Bright Field/Digital Phase Contrast Images)**: This section details how to generate a protocol that will allow for bright field kinetic imaging (converted to Digital Phase Contrast) in a single focal plane of view to determine PDO growth over time. The reason why users may choose to image in a single focal plane rather than generating a Z-Stack projection is because if the seeding density is too high, the PDOs overlap in different focal planes. This will then make it difficult for the analysis software to differentiate individual PDOs from each other.

1. Launch Gen5 software to begin imaging the 96-well plate.
2. **Click** New Task > Instrument Control > Incubate.
  1. Set the Requested temperature to 37 °C and **check** ON.
    1. *Note: Cytation will take a couple of minutes to reach temperature. Prior to placing the plate in the Cytation 5, make sure the reader is at 37 °C. This is necessary to maintain the sample at the appropriate temperature as well as decrease condensation on the lid, which will obstruct imaging*.
  2. **Close** the Instrument Control Panel.
3. Place plate in Cytation 5. **Click** New Task > Imager Manual Mode > Capture Now and input the following settings: Objective (select desired magnification); Filter (select microplate); Microplate format (select number of wells); and Vessel Type (select plate type). **Click** “Use Lid” and “Use slower carrier speed.” **Click** OK.
  1. *Note for Vessel Type: Be as specific as possible when selecting information about the plate because the software is calibrated to the specific distance from the objective to the bottom of the plate for each plate type as well as the thickness of the plastic*.
  2. *Note for Slower Carrier Speed: Select this box to avoid disrupting organoid domes when loading/unloading plates*.
4. Identify the focal plane.
  1. **Select** a well of interest to view (left panel, below histogram).
  2. **Select** the Bright Field channel (left panel, top).
  3. Use the coarse and fine adjustment arrows (left panel, middle) to change the focal plane in view.
    1. *Note: The distance at which each tick changes the focal height, for both coarse and fine adjustment, can be lowered or increased using the sliders under the Focus drop down menu*.
  4. Identify the bottom and top focal heights of the domes and choose the focal height that falls in the middle of these two values.
    1. *Note: For users using Agilent 96-Well Plates and seeding 5 µL domes, this focal height will be approximately 3700 µm*.
  5. To ensure that the focal height settings are appropriate for other wells of interest, **select** another well (left panel, below histogram) and visualize this focal height to make sure the image is still in focus. This is done by manually entering the focal positions. **Click** on the three dots next to the fine adjustment (left panel, top). A window will open. Type in the desired focal height.
5. Set the exposure settings for the Bright Field channel.
  1. First **Click** Auto Expose (left panel, top, under coarse and fine adjustment) to automatically determine an exposure that the Cytation 5 deems appropriate.
    1. If this exposure appears too dim or too bright, this can be adjusted manually using the plus and minus buttons on either side of the Auto Expose button.
6. Generate a template image from which the protocol/experiment will be based.
  1. **Click** on the “Camera” icon (left panel, bottom corner) to take a template image. This is what the images will look like when carrying out actual experiments.
  2. **Click** the “Process/Analyze” button to the right of the “Camera” icon.
  3. **Click** “Image Set” drop down menu (top left of the screen) and **click** on “Create experiment from this image set”. A new Procedure window will open.
    1. *Note: The parameters selected for the image will automatically be taken into the new window whereby an experimental protocol can be created*.
7. Create Protocol.
  1. Set the temperature and gradient: **Click** on Set Temperature under the Actions heading (left). A new window will open. **Select** “Incubator On” and manually enter the desired temperature under “Temperature.” Next, under “Gradient”, manually enter “1.” Close window by **selecting** OK.
    1. *Note: creating a 1 °C gradient will prevent condensation from forming on the lid of the plate*.
  2. Designating wells to image
    1. **Double click** on the Image tab under description.
    2. **Click** “Full Plate” (right corner, top). This will open the Plate Layout window.
    3. Highlight wells of interest using the cursor. **Click** OK.
    4. If desired, **check** “Autofocus binning” and “Capture binning” boxes. **Click** OK to close window.
      1. Note: Please refer to *Data Management* in the Discussion for specific scenarios in which this feature may be used.
  3. Set intervals for kinetic imaging.
    1. **Click** on Options under the Other heading (left).
    2. **Check** the “Discontinuous Kinetic Procedure” box.
    3. Under Estimated total time, enter the run time for the experiment (e.g., 5 days). Under Estimated interval, enter the interval at which to image the plate (e.g., every 6 hours).
    4. **Click** “Pause after each run” to allow time for the plate to be transferred to the BioSpa incubator. Close window by **selecting** OK.
  4. Set up Data Reduction to generate Digital Phase Contrast Images. Converting Bright field images into Digital Phase Contrast images allows users to more accurately create masks around objects of interest even when PDOs are undergoing cell death and blebbing, which can interfere with generation of masks around live/viable PDOs.
    1. In the toolbar, **click** Protocol > Data Reduction > Digital Phase Contrast, which will open a new tab.
    2. Make sure the “Channel” is set to Bright Field and set the “Structuring Element Size” to the average size at which PDOs are expected to grow. **Click** “OK” to close the window, and then **click** “OK” again to close the Data Reduction window.
  5. Save the Protocol.
    1. In the toolbar, **Click** File tab > Save Protocol as.
    2. **Select** location to save file. Enter file name. **Click** Save to close window.
    3. **Click** on the File > Exit in the toolbar. A tab will open to save changes to Imager Manual Mode. **Select** No.
    4. A tab will open to save changes to Experiment 1. **Select** No.
    5. A tab will open to update the protocol definition. **Select** Update.
    6. **Close** Gen5 software.
8. Import the Protocol into BioSpa OnDemand and finish setting up the Experiment.
  1. **Open** the BioSpa OnDemand software.
  2. **Select** an available slot in the BioSpa.
  3. Import the Protocol.
    1. Under the Procedure Info tab, **select** User in the drop-down menu.
    2. Next to Protocol slot, **click** Select > Add a new entry.
    3. Next to Protocol slot, **click** Select. This will open a new window to navigate to the desired Protocol in the file architecture.
    4. **Click** Open to import the Protocol into the BioSpa OnDemand software.
    5. Enter the amount of time needed to image the plate. **Click** OK to close the Gen5 Protocol List window.
      1. *Note: This step is especially important when running several experiments at a time. To determine the time needed to image the, click “Perform a timing run now*.*” Click OK*.
  4. Set imaging intervals and schedule the experiment.
    1. Under Interval, enter the imaging interval which was designated previously in *Step 5*.*4* of *Setting up Imaging Parameters*.
    2. Under Start Time Options, **select** “When available.”
      1. *Note: A specific start time can be designated instead of running the protocol at the next available time*.
    3. Under Duration, **select** “Fixed” or “Continuous.”
      1. *Note: Selecting Fixed duration will set a specific endpoint for the experiment and requires the user to designate an experimental timeframe. Continuous duration will allow the experiment to run with no endpoint and can only be ended by a user stopping the experiment*.
    4. **Click** Schedule plate/vessel. This will open the Plate Validation Sequence.
    5. A tab will open with the proposed first read time. **Click** Yes to accept this schedule.
  5. Remove the plate from the Cytation 5. **Click** Open drawer to access the appropriate slot. Place plate in BioSpa. **Click** Close drawer.
    1. *Note: This step can be performed at any point once the Protocol has been created. However, the plate must be in the Cytation 5 if one wishes to perform a timing run*.

**Digital Phase Contrast Image analysis in Gen5 software**: Below we describe methods to analyze data from the Digital Phase Contrast images generated from the Bright Field images. Representative images are provided in **Supplemental Figure S4**.

1. Opening image analysis module.
  1. **Open** experimental file in Gen5 software. **Select** Plate > View from the toolbar.
  2. **Change** Data drop-down menu to Dig.Ph.Con.
  3. **Double click** on a well of interest.
  4. **Select** Analyze > “I want to setup a new Image Analysis data reduction step” > OK.
2. Cellular Analysis.
  1. Primary Mask
    1. Under ANALYSIS SETTINGS, **select** Type: Cellular Analysis and Detection Channel: Dig.Ph.Con. (left panel, center).
    2. **Click** Options. The Primary Mask and Count page will open. In the Threshold box, **uncheck** “Auto” and adjust the slider as necessary to include or exclude objects of interest.
      1. *Note: When analyzing images in the Bright Field channel, ensure that Background is set to Light and for Digital Phase Contrast channel use Dark*.
      2. **Check** both boxes “Split touching objects” and “Fill holes in masks.”
      3. **Open** “Advanced Detection Options.”
      4. **Select** “Background Flattening” and “Auto.”
        1. *Note: The Rolling Ball Diameter is a pre-processing technique where the image is sampled to distinguish background noise from actual signal. The diameter is how much of the image is sampled*.
      5. **Set** “Image Smoothing Strength” to between 1 and 10 cycles of 3x3 average filter depending on how much background material there is.
        1. *Note: Image smoothing is used to further decrease the impact of background noise on the generation of the mask, it reduces the variability of background signal to allow for more accurate border identification and better special measurements*.
      6. **Set** the “Primary Mask” to “Use Threshold Mask” from the drop-down menu, then **select** “OK”.
      7. Under Object selection, designate a minimum and a maximum object size (µm). Adjust as necessary to exclude cellular debris/single cells.
        1. *Note: PDO size may vary significantly between different models and types. Use the measuring tool* 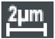 *in the Gen5 software to determine the minimum and maximum PDO size thresholds for each model*.
        2. **Deselect** “Include primary edge objects” and “Analyze entire image.” To limit the analysis to a certain region of the well, **click** “Plug.” This will open the Image Plug Window. Using the drop-down menu, **select** Plug shape and adjust size and position parameters to fit over the region of interest.
          1. *Note: It is important to maximize the number of PDOs within the plug while also excluding PDO-free areas to minimize background. Designate a plug size that will consistently capture the majority of the objects of interest across replicates. Generating a plug that also excludes the edges of the dome is important as it excludes any objects that may appear distorted due to the refraction of light from the extreme curvature of the dome around the edges*.
  2. Subpopulation Analysis.
    1. **Click** on Calculated Metrics in the Cellular Analysis toolbar. **Click** “Select or create object level metrics of interest” (right corner, bottom). Under Available object metrics, **select** metrics of interest (e.g., Circularity, StdDev) and **click** the Insert button. **Click** OK.
      1. *Note: Morphology and density of each PDO model will determine the best metrics of interest to distinguish the subpopulation; for analysis of Digital Phase Contrast images, circularity and StdDev are the typical metrics of use. Circularity allows for exclusion of cellular debris that do not have a more typical uniform circular structure. StdDev is distinguishes between cellular debris and PDOs. Specifically, debris will appear uniformly bright whereas PDOs will have brighter edges and darker cores and therefore a high StdDev of light*.
    2. **Open** the Subpopulation Analysis page from the Cellular Analysis toolbar. **Click** Add to create a new subpopulation. A pop-up window will open.
    3. If desired, enter a name for the subpopulation. Under Object metrics, **click** on metric of interest and **press** Add Condition. In the Edit Condition window, enter parameters for the chosen Object metric. Repeat with additional metrics as necessary.
      1. *Note: Parameters may be adjusted manually (i*.*e*., *include all objects with a circularity greater than 0*.*3)*.
    4. In the table at the bottom of the window, **check** the desired results to display. **Click** OK > Apply.
    5. To view the objects within the subpopulation, use the Object details drop-down menu to **select** the subpopulation. Objects that fall within the parameters will be highlighted in the image.
    6. To adjust subpopulation parameters, **reopen** the Subpopulation Analysis window from the Cellular Analysis toolbar. **Select** the subpopulation and **click** Edit.
    7. **Click** ADD STEP.
      1. *Note: This will apply the same analysis to all wells within the experiment at all time points. In the drop-down menu on the Matrix page, different metrics can be selected for individual viewing*.

**End of treatment cell viability and cell death fluorescence imaging using Nexcelom Bioscience ViaStain™ AOPI Staining Solution:** This section details the experimental procedure and parameters used to analyze cell viability and cell death within the organoid cultures using fluorescence. AOPI is a combination of two reagents, acridine orange (AO) and propidium iodide (PI). AO can enter both live and dead cells, resulting in the staining of all nucleated cells; AO generates a green fluorescent signal. PI can only enter cells with compromised membranes, resulting in staining all dead nucleated cells; PI generates a red fluorescence signal. Due to Förster resonance energy transfer (FRET), the PI signal quenches the AO signal in cells stained with both dyes, resulting in no spill-over and no double positive results.

1. Addition of “Viastain™ AOPI Staining Solution” to PDO culture.
  1. Add AOPI staining solution at a 1:50 v/v ratio (e.g., 2 µL of staining solution to 100 µL culture medium) to each well being careful not to introduce any air bubbles.
  2. Gently shake the plate to mix the AOPI solution with the culture medium and incubate in a dark place for 25 minutes before continuing.
    1. *Note: Future experiments should be incubated with the AOPI solution for 30 minutes before reading*.
2. To set up a new protocol for the AOPI analysis, repeat steps 1-6 from “Setting up imaging parameters for a single focal plane of view analysis (Bright Field/Digital Phase Contrast Images).”
3. Create Protocol.
  1. Set the temperature and gradient: **Click** on Set Temperature under the Actions heading (left). A new window will open. **Select** “Incubator On” and manually enter the desired temperature under “Temperature.” Next, under “Gradient”, manually enter “1.” Close window by **selecting** OK.
    1. *Note: creating a 1 °C gradient will prevent condensation from forming on the lid of the plate*.
  2. Formatting the plate layout and read description.
    1. **Double click** on the Image tab under Description.
    2. **Click** “Full Plate” (right corner, top). This will open the Plate Layout window.
    3. Highlight wells of interest that you wish to image using the cursor. **Click** OK.
    4. Under the image drop down menu (top, middle) **select** “Crop 75%.”
      1. *Note: Selecting the “crop 75%” option reduces the amount of background fluorescence that will naturally occur around the edges of the wells as the field of view being imaged is slightly decreased*.
    5. **Select** both “Autofocus binning” and “Capture binning.”
    6. Under “Channels” there should be one current channel selected “Bright Field;” **select** the number “2” to add a second channel.
      1. Under the Color drop down menu **select** “GFP 469,525.”
      2. **De-select** “Auto” and then **click** on the microscope icon next to “Auto.”
        1. *Note: This step will allow for manual setting of the exposure settings*.
      3. Adjust the “Illumination intensity,” “Integration time” and “Camera gain” to appropriate values so that exposure levels are correct.
    7. Repeat the previous step (3.2.6) but instead **select** the number “3” to add a third channel, and under the color drop down menu **select** “Texas Red 586,647.”
    8. Once the three channels are set up, **click** “OK” to close the “Imaging Step-Inverted Imager” tab.
  3. **Click** “Validate” at the bottom of the “Procedure” window to confirm the procedure step sequence is valid and then **click** “OK” and then “OK” again.
    1. *Note: Raw images that are generated through this protocol will naturally have a lot of background fluorescence and therefore a Data Reduction>Image Preprocessing step needs to be implemented to normalize for background fluorescence*.
4. **Click** on the “Protocol” tab (top left) and **select** “Data Reduction.”
  1. Under “Image processing” **select** “Image Preprocessing.” A new window will open.
    1. *Note: The bright field image will not need any image preprocessing steps applied*.
  2. **De-select** “Apply image preprocessing” for the Bright Field channel.
  3. **Click** on the “GFP 469,525” tab.
    1. Make sure “Apply image preprocessing” is selected.
    2. **Select** “Dark” from the Background drop down menu.
    3. **De-select** “Use same options as channel 1.”
    4. Make sure “Background Flattening” and “Auto” is selected.
    5. Change the “Image smoothing strength” to 1 Cycle of 3x3 average filter.
  4. **Click** on the “Texas Red 586,647” tab and repeat steps 4.3.1-5.
  5. **Click** “OK” and then **click** “OK” again.
5. Save the protocol for future use by **clicking** on the “File” tab, top left of the screen, and then “Save Protocol As…”
  1. Name the protocol appropriately and **click** “Save.”

**End of treatment cell viability and cell death fluorescence image analysis in Gen5 software**: Below we describe methods to analyze data from the End of Treatment AOPI Fluorescence Protocol. Two separate image analysis steps need to be set up: 1) GFP channel, which is a measure of viability (AO); 2) Texas Red channel, which is the measure of cell death (PI).

1. Opening image analysis module.
  1. **Open** experimental file in Gen5 software. **Select** Plate > View from the toolbar.
  2. **Change** Data drop-down menu to Picture [Tsf[Bright Field+GFP 469,525+Texas Red 586,647]].
  3. **Double click** on a well of interest.
  4. **Select** Analyze > “I want to setup a new Image Analysis data reduction step” > OK.
2. Cellular Analysis for Cell Viability (Acridine Orange and GFP fluorescent channel).
  1. GFP Primary Mask
    1. Under ANALYSIS SETTINGS, **select** Type: Cellular Analysis and Detection Channel: Tsf[GFP 469,525] (left panel, center).
    2. **Click** Options. The Primary Mask and Count page will open. In the Threshold box, **check** “Auto” and adjust the slider as necessary to include or exclude objects of interest.
      1. *Note: When analyzing images using the GFP or Texas Red channels, set the background to dark*.
  2. **Select** “Split touching objects” and “Fill holes in masks.”
  3. **Open** “Advanced Detection Options.”
    1. **Select** “Background Flattening” and **de-select** “Auto.”
      1. *Note: The Rolling Ball Diameter is a pre-processing technique where the image is sampled to distinguish background noise from actual signal. The diameter should be set to roughly the size of the largest object being analyzed*.
    2. **Set** “Image Smoothing Strength” to 1 cycle of 3x3 average filter.
    3. **Set** the “Primary Mask” to “Use Threshold Mask” from the drop-down menu, and then **select** “OK.”
    4. Under Object selection, designate a minimum and maximum object size (µm). Adjust as necessary to exclude cellular debris/single cells.
      1. *Note: PDO size may vary significantly between different models and types. Use the measuring tool to determine the minimum and maximum PDO size thresholds for each model*.
      2. **Deselect** “Include primary edge objects” and “Analyze entire image.” To limit the analysis to a certain region of the well, **click** “Plug.” This will open the Image Plug Window. Using the drop-down menu, **select** Plug shape and adjust size and position parameters to fit over the region of interest.
      3. *Note: It is important to maximize the number of PDOs within the plug while also excluding PDO-free areas to minimize background. Designate a plug size that will consistently capture the majority of the objects of interest across replicates. Generating a plug that also excludes the edges of the dome is important as it excludes any objects that may appear distorted due to the refraction of light from the extreme curvature of the dome around the edges*.
  4. Subpopulation Analysis.
    1. **Click** on Calculated Metrics in the Cellular Analysis toolbar. **Click** “Select or create object level metrics of interest” (right corner, bottom). Under Available object metrics, **select** metrics of interest (e.g., Circularity, Integral[Tsf[GFP 469,525]]) and **click** the Insert button. **Click** OK.
      1. *Note: Morphology and density of each PDO model will determine the best metrics of interest to distinguish the subpopulation. For analysis of the GFP channel, circularity is the only metric required as only viable material will fluoresce green and therefore there is no need for exclusion of debris*.
    2. **Open** the Subpopulation Analysis page from the Cellular Analysis toolbar. **Click** Add to create a new subpopulation. A pop-up window will open.
    3. If desired, enter a name for the subpopulation. Under Object metrics, **click** on metric of interest and **select** Add Condition. In the Edit Condition window, enter parameters for the chosen Object metric. Repeat with additional metrics as necessary.
      1. *Note: Parameters may be adjusted manually (i*.*e*., *include all objects with a circularity greater than 0*.*3)*.
    4. In the table at the bottom of the window, **select** the desired results to display. **Click** OK > Apply.
    5. To view the objects within the subpopulation, use the Object details drop-down menu to **select** the subpopulation. Objects that fall within the parameters will be highlighted in the image.
    6. To adjust subpopulation parameters, **reopen** the Subpopulation Analysis window from the Cellular Analysis toolbar. **Select** the subpopulation and **click** Edit.
    7. **Click** ADD STEP.
      1. *Note: This will apply the same analysis to all wells within the experiment at all time points. In the drop-down menu on the Matrix page, different metrics can be selected for individual viewing*.
3. Cellular Analysis for Cell Death (Propidium Iodide and Texas Red fluorescent channel).
  1. Texas Red Primary Mask.
    1. Under ANALYSIS SETTINGS, **select** Type: Cellular Analysis and Detection Channel: Tsf[Texas Red 586,647] (left panel, center).
    2. **Click** Options. The Primary Mask and Count page will open. In the Threshold box, **uncheck** “Auto” and set the value to 5000.
      1. *Note: When analyzing images using the GFP or Texas Red channels, set the background to dark*.
  2. **Check** both boxes “Split touching objects” and “Fill holes in masks.”
  3. **Open** “Advanced Detection Options.”
    1. **Select** “Background Flattening” and **select** “Auto.”
    2. **Set** “Image Smoothing Strength” to 0 cycles of 3x3 average filter.
    3. **Set** the “Evaluate background on” to 85% of lowest pixels.
      1. *Note: This step is needed to ensure that no background specks of fluorescence are include in the mask for analysis*.
    4. **Set** the “Primary Mask” to “Use Threshold Mask” from the drop-down menu, then **select** “OK.”
      1. *Note: For the Texas Red channel, minimal image preprocessing techniques are needed for the analysis since the fluorescence is more focal*.
  4. Under Object selection, designate a minimum and a maximum object size (µm).
    1. *Note: For the Texas Red channel analysis, the minimal object size should be approximately the size of one cell (e*.*g*., *10 µm), but the maximum should still be that of the largest object expected for the given model*.
  5. **Deselect** “Include primary edge objects” and “Analyze entire image.” To limit the analysis to a certain region of the well, **click** “Plug.” This will open the Image Plug Window. Using the drop-down menu, **select** Plug shape and adjust size and position parameters to fit over the region of interest. This should be the same size and position as the plug used for the GFP channel.
    1. *Note: It is important to maximize the number of PDOs within the plug while also excluding PDO-free areas to minimize background. Designate a plug size that will consistently capture the majority of the objects of interest across replicates. Generating a plug that also excludes the edges of the dome is important as it excludes any objects that may appear distorted due to the refraction of light from the extreme curvature of the dome around the edges*.
  6. **Click** ADD STEP.
    1. *Note: This will apply the same analysis to all wells within the experiment at all time points. In the drop-down menu on the Matrix page, different metrics can be selected for individual viewing*.
    2. *Note: There is no need for a subpopulation analysis for the Texas Red channel as the majority of the signal will be focal and will stem from dead material. Therefore, this signal should not be excluded from analysis*.

---

**Supplementary Table S1:** Organoid Culture Media Components. Note that reagents have been optimized for culturing gynecologic cancer PDOs.

| Organoid Culture Media Component | Final Concentration |
| --- | --- |
| AdDF+++ (Advanced DMEM/F12, Glutamax, HEPES, penicillin/streptomycin) | 1X Glutamax, 10 mM HEPES, 10 units/mL penicillin/streptomycin |
| Noggin | 100 ng/mL |
| Respondin-1 | 250 ng/mL |
| A83-01 | 500 nM |
| FGF-10 | 10 ng/mL |
| Heregulinβ-1 | 37.5 ng/mL |
| EGF | 0.5 ng/mL |
| Estradiol | 100 nM |
| Y-27632 | 5 μM |
| Forskolin | 10 μM |
| Hydrocortisone | 500 ng/mL |
| B27 supplement | 1X |
| Nicotinamide | 10 mM |
| N-acetylcysteine | 1.25 mM |
| Primocin | 100 μg/mL |

**Supplementary Figure S1:**
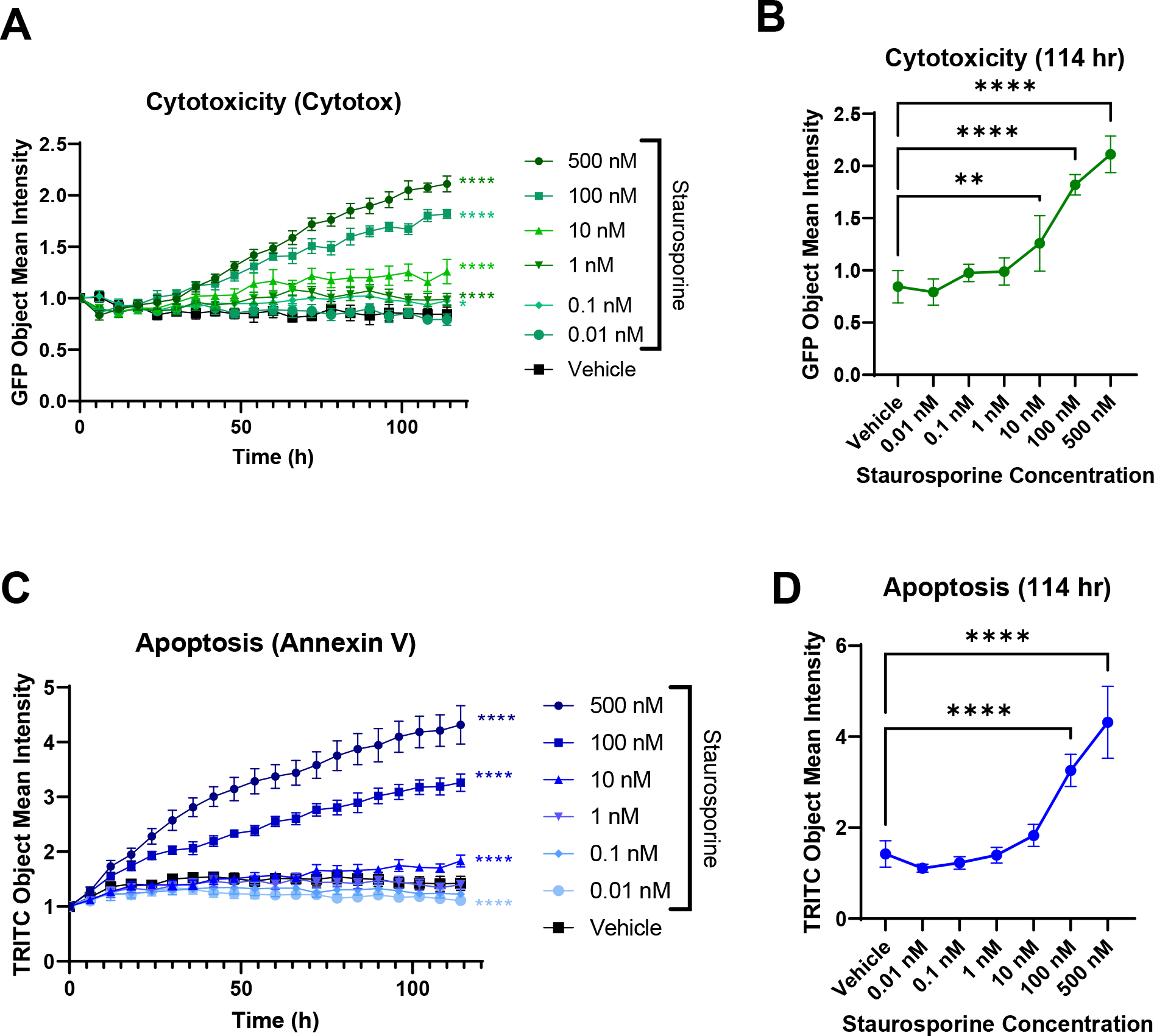
Multiplexed live-cell imaging of ONC-10811. PDOs were plated in 96-well plates and incubated in Annexin V Red (1:400) and Cytotox Green (200 nM) dyes overnight at 37 °C. The following day, PDOs were treated with increasing concentrations of staurosporine and were imaged every 6 hours for 5 days. (**A**) Time and dose-dependent increase in cytotoxicity in response to staurosporine. Data were plotted as the Object Mean Intensity in the GFP channel. (**B**) Dose-response of staurosporine at 114 hrs. Data were plotted as the Object Mean Intensity values in the GFP channel at the 114 hr timepoint. (**C**) Time and dose-dependent increase in apoptosis in response to staurosporine. Data were plotted as the Object Mean Intensity in the TRITC channel. (**D**) Dose-response of staurosporine at 114 hrs. Data were plotted as the Object Mean Intensity values in the TRITC channel at the 114 hr timepoint. Data in A and C were normalized to PDO number at time 0 at the well level. N=5 technical replicates per treatment in each model. **** p < 0.0001 vs. vehicle control by 2-way ANOVA.

**Supplementary Figure S2:**
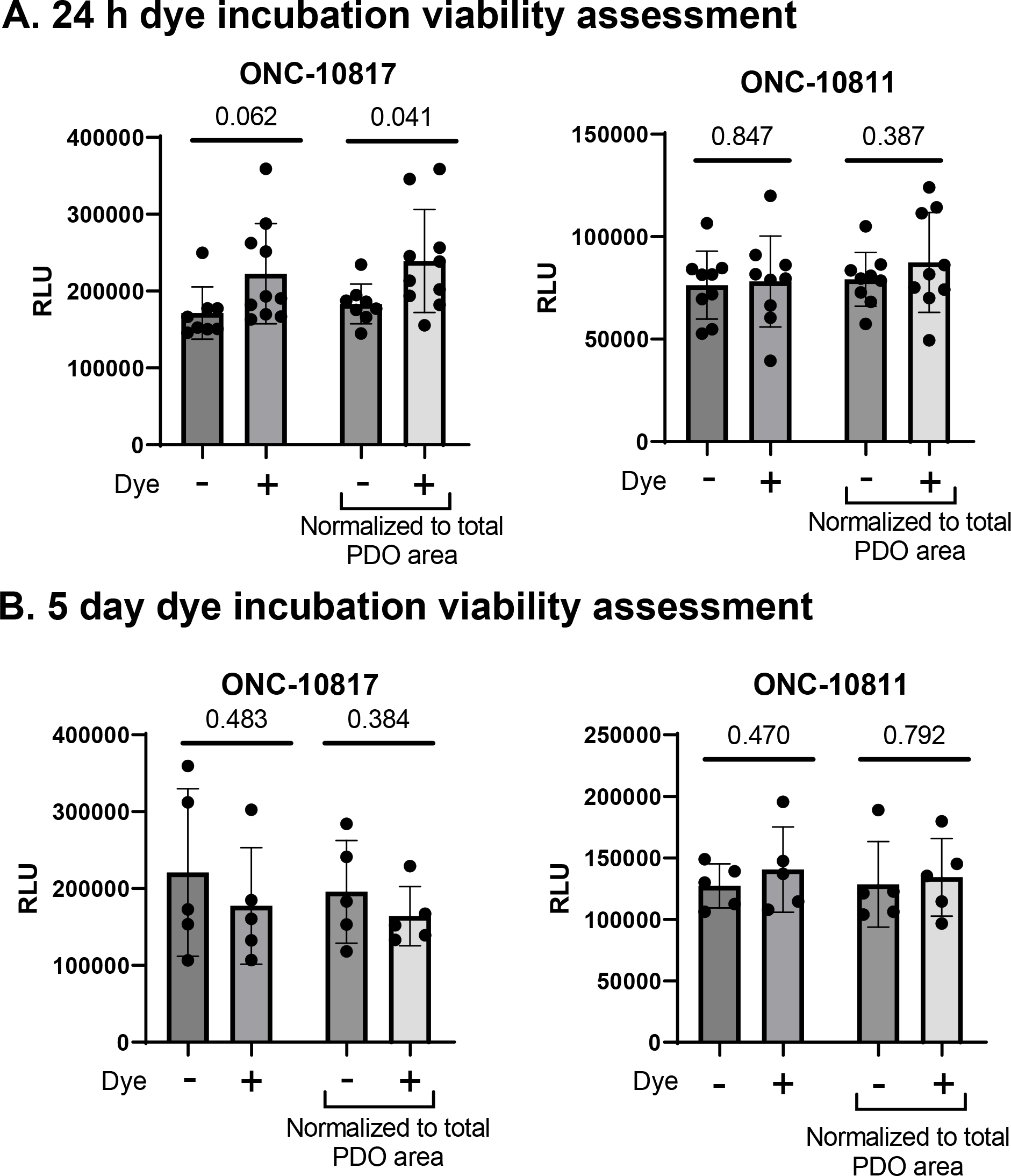
Treatment with Annexin V and Cytotox does not perturb PDO viability. PDOs were plated in 96-well plates and incubated with Annexin V Red (1:400) and Cytotox Green (200 nM) dyes overnight at 37 °C. Following the 24 hour incubation, viability was assessed using the CellTiter-Glo 3D assay per the manufacturer’s protocol. (**A**) Viability in dye-treated and untreated PDOs at the 24 hr timepoint. Both relative light unit (RLU) values and values normalized to organoid sum area are presented. Specifically, the CellTiter-Glo3D RLU for each well was normalized to the summation of organoid area for that well at the time of plating (i.e., immediately after dye addition). Total PDO area was determined using the “Sum Area” calculation in Cellular Analysis. N=10.(**B**) Viability in dye-treated and untreated PDOs at the 114 hr timepoint. Both raw luminescence values and values normalized to total PDO area are presented. The CellTiter-Glo3D RLU for each well was normalized to the summation of organoid area at 24 hours post-plating, which corresponds to time 0 in the kinetic imaging experiments. N=5. Significance in A and B was assessed using an unpaired t-test; p values are listed on the graphs.

**Supplementary Figure S3:**
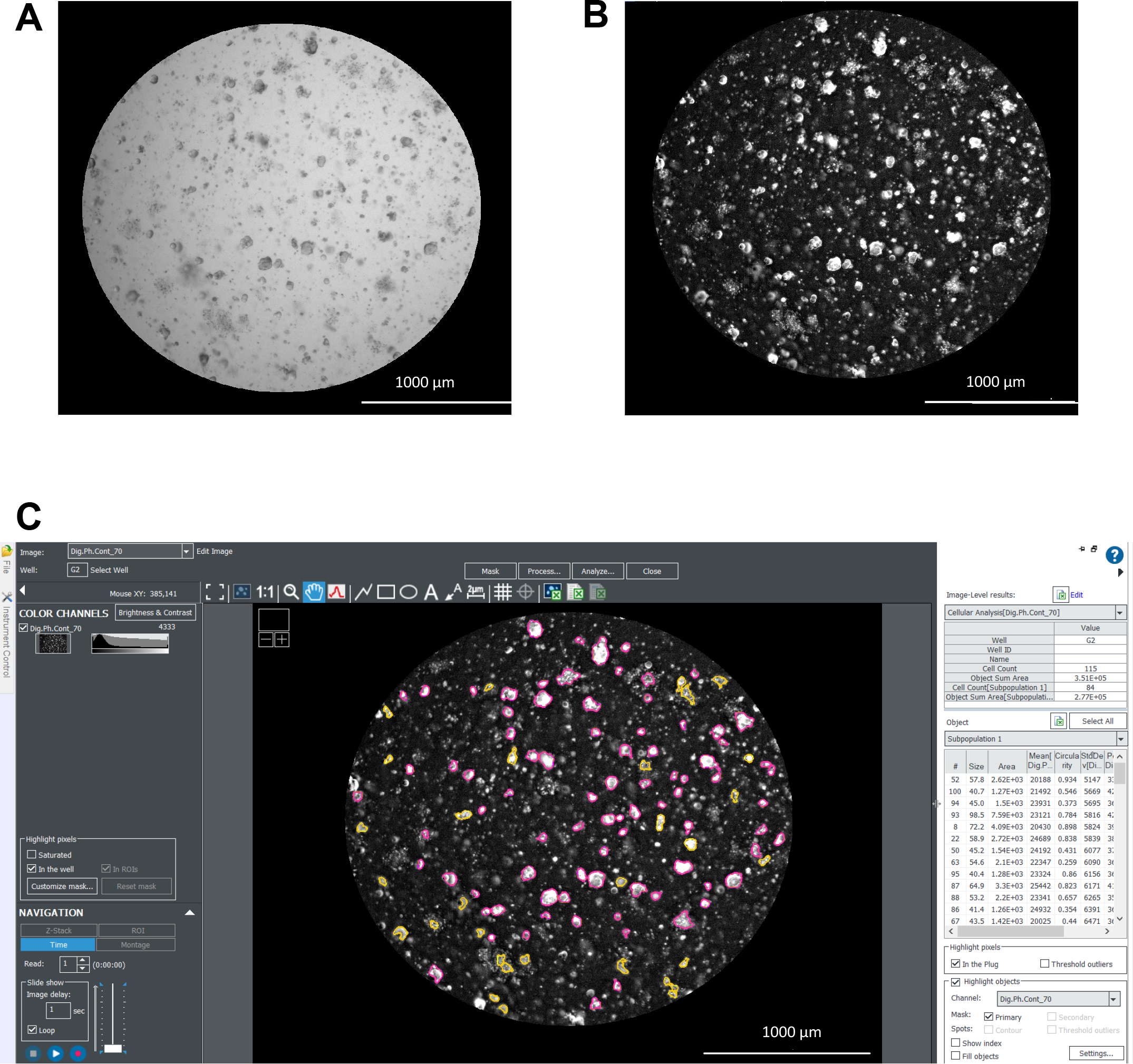
Label-free analysis of PDOs using digital phase contrast. This figure contains representative images (2.5X objective) depicting label-free analysis as described in the **Supplemental Protocols:** *Setting up imaging parameters for a single focal plane of view analysis (Bright Field/Digital Phase Contrast Images) and Digital Phase Contrast Image analysis in Gen5 software*. (**A**) Example Bright Field image of a prostate cancer PDXO model at a single focal plane. (**B**) The Bright Field image in **A** was converted to a Digital Phase Contrast image. Dark objects in the Bright Field image appear bright in the Digital Phase Contrast image and vice versa. (**C**) Example of PDO masking using the Digital Phase contrast image. Note that the edges of objects of interest become much more defined as compared to the Bright Field images. In this representative image, objects in the primary mask are outlined in yellow. The subpopulation in (C) (outlined in pink) is defined based on circularity (>0.3), area (>1000), Mean[Dig.Ph.Con] > 2000, StdDev[Dig.Ph.con] > 5000 + < 13500, and Peak[Dig.Ph.Con] > 12500. The parameters used in this representative figure were applied to data in Figure 8 to normalize to cell count on day 0.

**Supplementary Figure S4:**
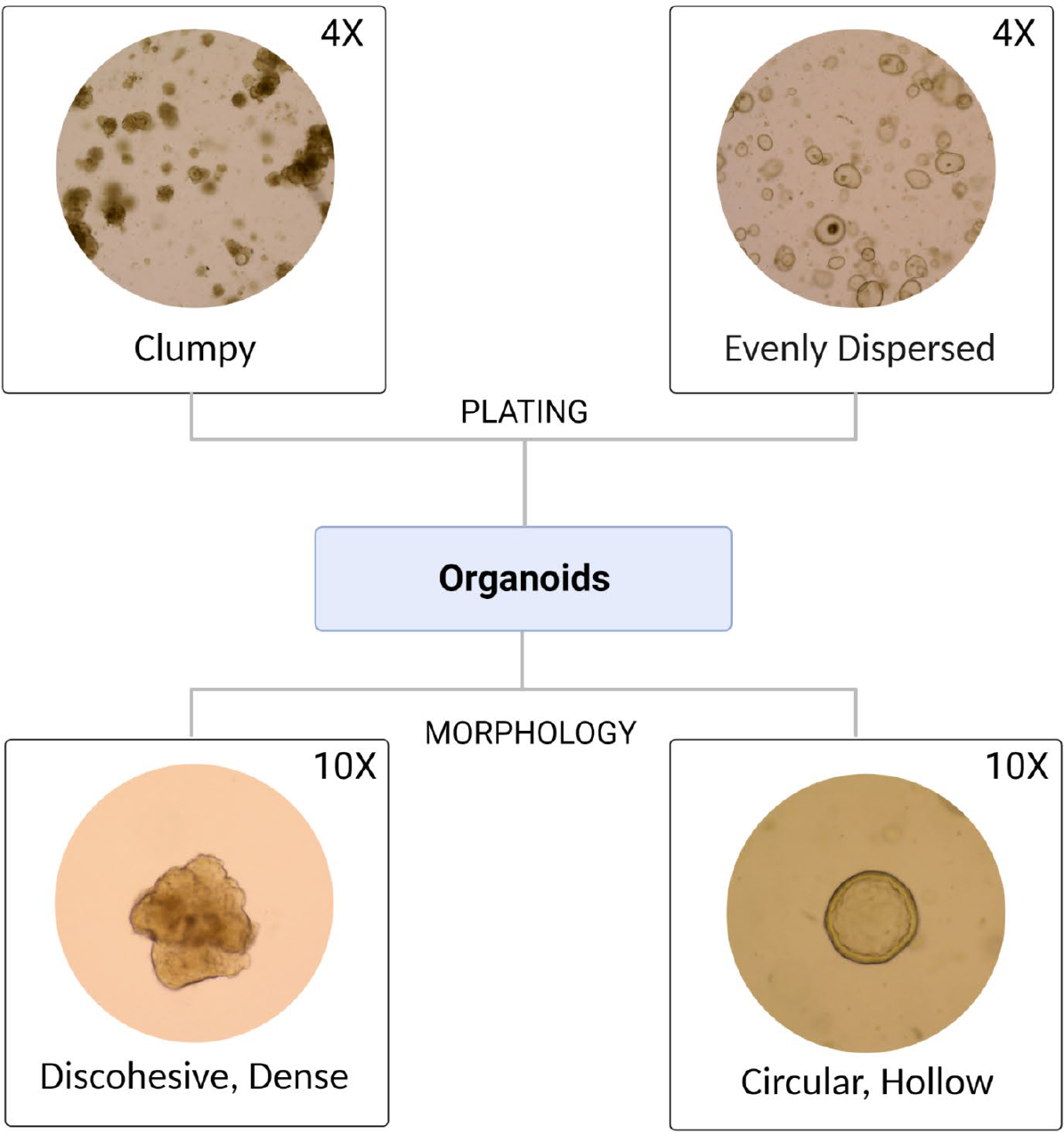
PDO models may vary in their morphology and plating consistency. *Upper Panel*: Examples of differential dispersion of PDOs in the BME domes. *Lower Panel:* Representative images of discohesive vs. circular PDOs. All images were acquired with an EVOS microscope. Magnifications are noted.

**Supplementary Figure S5:**
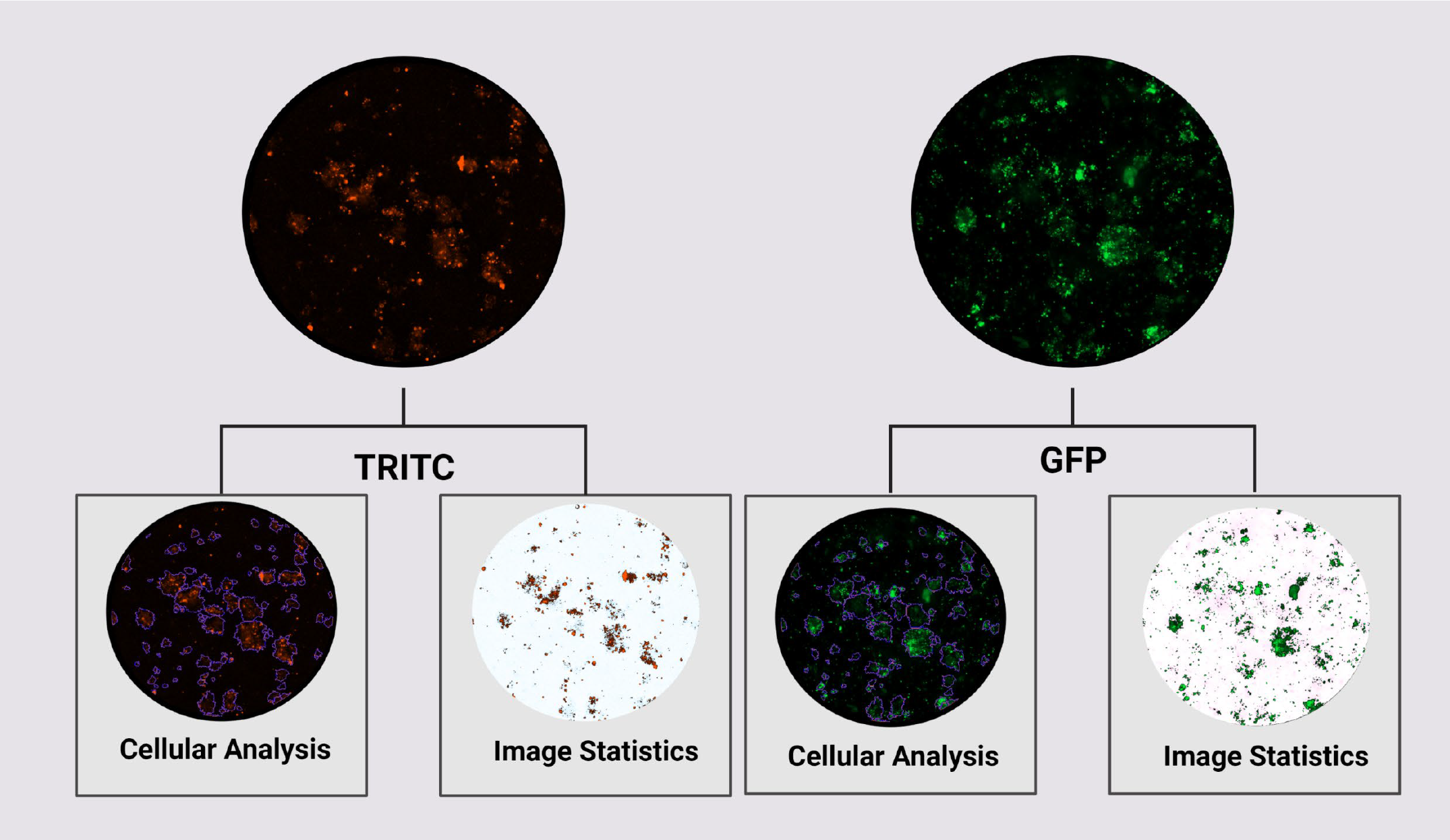
Example images using Cellular Analysis vs. Image Statistics for quantifying fluorescence. Using Cellular Analysis, users can define specific populations within an image and measure fluorescence in those regions. Image Statistics may also be used to measure fluorescence in an image by defining a threshold to exclude background signal. *See Discussion for the limitations of using Image Statistics*. Images are at 4X magnification.

